# The DNA Damage Response (DDR) landscape of endometrial cancer defines discrete disease subtypes and reveals therapeutic opportunities

**DOI:** 10.1101/2023.11.20.567919

**Authors:** Xingyuan Zhang, Sayali Joseph, Di Wu, Jessica L. Bowser, Cyrus Vaziri

## Abstract

Genome maintenance is an enabling characteristic that allows neoplastic cells to tolerate the inherent stresses of tumorigenesis and evade therapy-induced genotoxicity. Neoplastic cells also deploy mis-expressed germ cell proteins termed Cancer Testes Antigens (CTAs) to promote genome maintenance and survival. Here, we present the first comprehensive characterization of the DNA Damage Response (DDR) and CTA transcriptional landscapes of endometrial cancer in relation to conventional histological and molecular subtypes. We show endometrial serous carcinoma (ESC), an aggressive endometrial cancer subtype, is defined by gene expression signatures comprising members of the Replication Fork Protection Complex (RFPC) and Fanconi Anemia (FA) pathway and CTAs with mitotic functions. DDR and CTA-based profiling also defines a subset of highly aggressive endometrioid endometrial carcinomas (EEC) with poor clinical outcomes that share similar profiles to ESC yet have distinct characteristics based on conventional histological and genomic features. Using an unbiased CRISPR-based genetic screen and a candidate gene approach, we confirm that DDR and CTA genes that constitute the ESC and related EEC gene signatures are required for proliferation and therapy-resistance of cultured endometrial cancer cells. Our study validates the use of DDR and CTA-based tumor classifiers and reveals new vulnerabilities of aggressive endometrial cancer where none currently exist.

## Introduction

Genome maintenance is an important enabling characteristic of cancer that confers tolerance of intrinsic and therapy-induced DNA damage. Different cancers arise from different precursors, are driven by different oncogenes, and experience different genotoxic exposures. Therefore, each cancer relies on distinct genome maintenance mechanisms to survive the unique stresses experienced during tumorigenesis. Because genome maintenance mechanisms can be error-free or error-prone, the availability and choice of pathways deployed by cancer cells also shapes the mutational landscape. Elucidating the genome maintenance pathways that sustain each cancer reveals mechanistic underpinnings of tumorigenesis. Moreover, the genome maintenance-dependencies of cancer cells are molecular vulnerabilities and may provide opportunities for therapy.

To define the genome maintenance processes that sustain cancer cells, we recently developed predictive classifiers based upon the relative expression levels of genes encoding core components of the major DNA Damage Response (DDR) and DNA repair pathways.^1^ In an initial study, our unsupervised analysis of DDR gene expression patterns in breast cancer patients (from both TCGA and North Carolina Breast Cancer cohorts) identified 4 distinct patient groups and, remarkably, predicted disease subtypes with accuracy.^1^ Tumor stratification based on DNA repair profiling is a highly innovative approach that reveals underlying mechanisms of carcinogenesis and is highly actionable. The repertoire of DDR pathways available to any tumor has the potential to determine responsiveness to chemotherapy. Therefore, DDR-based classifiers could be used to accurately predict sensitivities to distinct genotoxic therapeutics and inform clinical decisions. By analogy, the PAM50 signature helps guide clinical strategy in breast cancer.^2,3^

Here, we have used DDR-based transcriptional profiling to define the genome maintenance landscape of endometrial cancer (EC). Endometrial cancer is the only major solid tumor for which incidence and mortality rates are rising, underscoring the need for improved strategies to predict, prevent, and treat the disease. The DDR landscape of different endometrial cancer subtypes, and the contribution of the DDR to tumorigenic phenotypes in this cancer setting have not been studied.

Endometrial tumors are commonly (∼80%) of the endometrioid histotype (designated endometrial endometrioid carcinomas or EEC). There is tremendous heterogeneity in tumors belonging to the EEC subtype. EEC comprise both low grade (‘G1’ and ‘G2’), largely indolent tumors, and high grade (‘G3’) tumors that are a combination of indolent and highly aggressive tumors. Endometrial Serous Carcinomas (ESC) represent a distinct, highly aggressive, and historically uncommon subtype of endometrial tumors. ESC comprise ∼10% of all endometrial tumors yet account for ∼40% of all endometrial cancer deaths.^4^ The ongoing increase in endometrial cancer mortality is due to a disproportionate rise in ESC.^5^

In addition to histology-based subtyping, a comprehensive study of EEC and ESC by The Cancer Genome Atlas (TCGA) revealed four genomic subtypes of endometrial cancer.^6^ The *POLE* subtype with endometrioid histology is the least common, is ‘ultra-mutated’, and typically has good prognosis. The Microsatellite Instability (MSI) subtype also contains ‘hypermutated’ tumors of endometrioid histology. The copy-number variation (CNV)-low subtype comprises the majority of low grade endometrioid tumors. The CNV-high subtype comprises the most aggressive endometrioid tumors and all of ESC.^6^ Subsequent proteogenomic studies have revealed differences in underlying pathways and the immune landscapes of the TCGA subtypes.^7^

A defining feature of ESC (and the aggressive CNV-high endometrioid tumors) is their exceptional resistance to DNA-damaging therapies (e.g. carboplatin)^8^. Unfortunately, inherent or acquired DNA damage tolerance (‘chemoresistance’) occurs in 90% of ESC and patients with metastatic or recurrent EEC and is the major cause of death. Current gaps in knowledge of how ESCs and aggressive EECs tolerate chemotherapy limit development of life-saving treatment strategies. To improve patient outcomes, it is critical to elucidate how endometrial cancers tolerate therapy and become chemoresistant. Accordingly, we have sought to identify unique underlying features of aggressive endometrial cancers that account for their chemoresistance.

Our rationale is that a knowledge of the DDR landscape may reveal molecular underpinnings of chemoresistance and provide new opportunities for precision medicine. Our published DDR classifier gene set^1^ also included two representative Cancer/Testes Antigens (CTAs) with known roles in DNA repair. CTA genes are normally expressed only in the adult testes. However, many cancer cells aberrantly mis-express CTAs at high levels.^9^ CTAs often promote DNA repair, thereby conferring cancer cell fitness and tumorigenic phenotypes. For example, in lung cancer cells, Melanoma Antigen A4 (MAGE-A4) associates with the E3 ubiquitin ligase RAD18 and promotes post-replicative DNA repair.^10,11^ In lung cancer and breast cancer cell lines, HORMAD1 promotes Homologous Recombination (HR) and radio-resistance.^12,13^ Interestingly, expression levels of MAGEA4^10^ and HORMAD1^12,13^ were both highly associated with a specific breast cancer subtype.^1^ DDR-related functions have also been attributed to many additional CTAs.^14^ Owing to their absence from all normal somatic cells, CTA that promote DNA repair (and other tumorigenic phenotypes) represent very attractive therapeutic targets. Ablation of CTAs leads to reduced DNA repair capacity and chemo/radio-sensitivity yet is innocuous to non-cancerous cells.

In this study we have refined our methods for annotating tumors based upon DDR gene expression. Because CTAs provide a critical layer of regulatory input into DNA repair pathways in cancer cells^14^, we also interrogated CTA expression profiles in endometrial cancers. Using the TCGA uterine cancer dataset we demonstrate that the different subtypes of endometrial cancer are associated with distinct DDR and CTA expression profiles. Using independent approaches (including unbiased genetic screens) we validate the critical roles of DDR and CTA genes in sustaining growth of endometrial cancer cells. The gene signatures we derived are prognostic and provide new vulnerabilities that can be further harnessed for therapeutic benefit of endometrial cancer patients.

## Results

### DDR gene expression profiling reveals distinct endometrial cancer subsets

We curated a list of 131 genes encoding core components of 13 major DNA repair and genome maintenance pathways (Supplemental Table 1). Using the TCGA uterine cancer dataset we generated heatmaps depicting relative expression of these DDR genes in ESC and EEC. We determined relative expression levels of DDR genes in relation to multiple classifiers including demographic features (race and BMI), tumor grade, disease stage, and genome maintenance-related features (e.g., Replication Stress or RS signature; Homologous Recombination-Deficiency or HRD score; Centrosome Amplification or CA20 signature).

Histology, tumor grade, and disease stage are important independent prognostic factors for endometrial cancer.^15^ ESCs are often metastatic at time of diagnosis and have poor outcomes compared to more indolent endometrial tumors (G1 and G2 EEC). G3 EECs are heterogenous in that these tumors can be either indolent or aggressive and share similar molecular / genetic features with G1 and G2 EEC or ESC (e.g., *TP53* mutation). Race and BMI are also predictive factors. Lower BMI has been associated with higher disease stage and features of ESC (e.g., high HER2 expression),^16^ and Non-Hispanic Black women are more likely to be diagnosed with ESC and die of the disease compared to White women [https://seer.cancer.gov/statfacts/html/corp.html]. Oncogene-induced DNA replication stress is a major source of intrinsic DNA damage in cancer cells,^17^ which likely necessitates expression of DDR genes for stress tolerance. Different oncogenes induce different types of DNA damage and may create dependencies on distinct DDR pathways for damage tolerance.^18–20^ Therefore, in our classifiers we included various individual oncogenes / tumor suppressors that are frequently mutated or amplified in ESC and EEC (Fig 1A). Homologous Recombination Deficiency (HRD) is common in many tumors and creates new dependencies on alternative DDR mechanisms.^21^ Therefore, we considered that HR status may impact the expression and deployment of other DDR mediators in endometrial cancer. Because aberrant centrosome number is a potential source of mitotic errors and DNA damage in many tumors^22^, we also considered the Centrosome Amplification 20 (CA20) gene signature^23,24^ as a feature that might influence the DDR.

**Fig. 1.**
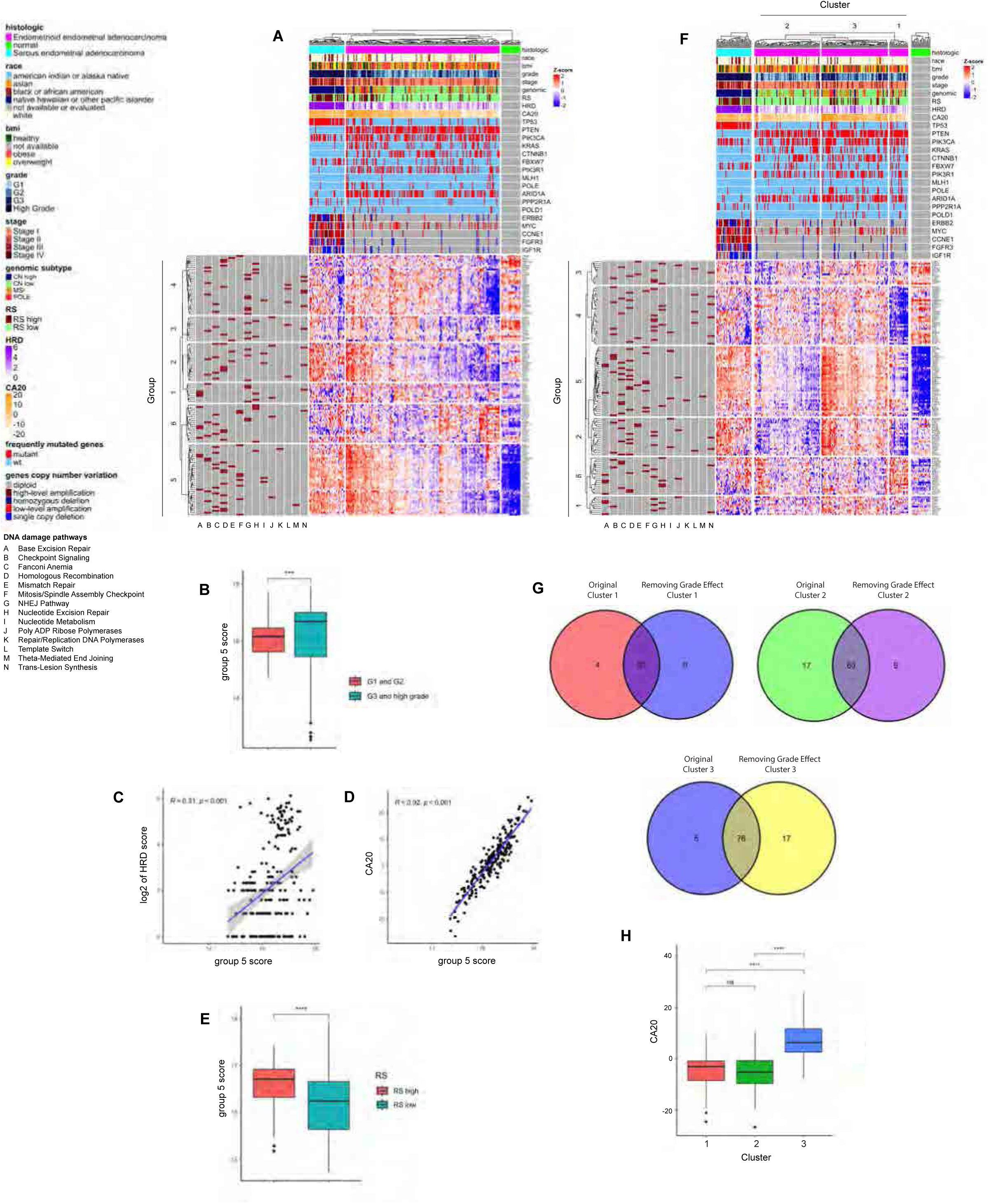
Expression of DDR genes in endometrial cancer with supervised clustering based on histological subtypes. (A) Genomic and clinical data from the 2013 TCGA sample set for uterine cancer were downloaded from TCGA portal and Xena Browser. Samples were clustered by endometrial tumor histology (serous or endometrioid) or as normal endometrium. Heatmaps are depicting relative mRNA expression of DDR genes assembled from a curated list of 131 genes that encode core components of 13 major DDR and genome maintenance pathways. Based on DDR gene expression patterns, K-means clustering (K = 6) identified 6 transcript gene signature groups (Groups 1-6). TransSig(Group 5) is highly expressed in serous endometrial tumors compared to normal endometrial tissue and contains 37 genes of different DDR pathways. DDR pathways and related genes are highlighted on the far left. Clinical, molecular, and genomic classifiers relevant to endometrial cancer and genomic maintenance features (HRD and CA20 scores) are depicted. Correlation of TransSig(Group 5) genes of endometrial serous tumors to tumor grade (B) and HRD (C) and CA20 (D) scores and replication stress (RS). TransSig(Group 5) gene score is the average expression of the genes in the cluster. (B) Wilcoxon’s rank sum test was applied to compare TransSig(Group 5) score with tumor grade (’G1’, ’G2’ as low grade, ’G3’ and high grade). (C-E) Spearman’s rank correlation. *p < 0.05, **p < 0.01, ***p < 0.001. (F) K-means clustering separating endometrioid tumors into 3 sub-clusters based on DDR gene expression profiles. (G) Venn diagrams depicting similar composition of the 3 sub-clusters for endometrioid tumors before (A) and after (Supplemental Fig. 2) removing tumor grade effect. (H) CA20 scores between the 3 sub-clusters for endometrioid tumors. (H) Wilcoxon’s rank sum test, ****p < 0.0001.

We performed supervised clustering and examined DDR gene expression patterns in relation to histological features of the tumors using the 2013 TCGA sample set for uterine cancer (n = 241).^6^ Based on their patterns of DDR gene expression, EEC and ESC tumors were both very distinct from normal tissue (Fig. 1A). ESCs are among the most chemoresistant and clinically challenging endometrial tumors. Therefore, it was of particular interest to define DDR features of ESC. Using K-means clustering, we identified 6 transcriptome gene signature groups of DDR genes, of which ‘transcriptome gene signature Group 2 (TransSigGroup2)’ and ‘transcriptome gene signature Group 5 (TransSigGroup5)’ were expressed at particularly high levels in ESC and a subset of EEC when compared with normal endometrial tissue (Fig 1A). DDR genes and associated pathways represented in TransSigGroup5 and TransSigGroup2 are listed in Table 1. We validated the differential expression patterns of TransSigGroup5 and TransSigGroup2 DDR genes using a secondary cohort that included all serous and endometrioid tumors in the TCGA database for uterine cancer (n = 521) (Supplemental Fig. 1). The DDR genes in TransSigGroup2 and TransSigGroup5 encode subunits of Ribonucleotide Reductase (RNR) and multiple genes involved in DNA replication fork progression. Most strikingly, TransSigGroup5 and TransSigGroup2 gene lists are characterized by transcripts encoding numerous mediators of the Fanconi Anemia (FA) and Homologous Recombination (HR) pathways (Table 1).

For both serous and endometrioid tumors, expression of TransSigGroup5 DDR genes associates with higher tumor grade (Fig. 1B), high HRD and CA20 scores (Fig. 1C-1D), and high RS (Fig. 1E).

In contrast with ESC, which showed a homogenous expression pattern for DDR genes in TransSigGroup5 and 2, expression of TransSigGroup5 and 2 genes was inherently heterogenous for EEC (Fig. 1A and Supplemental Fig. 1). Therefore, we used K-means clustering to further subcategorize endometrioid tumors into three distinct ‘Clusters’ based on their DDR gene expression patterns (Fig. 1F). The resulting Cluster 1 and Cluster 2 tumors expressed TransSigGroup5 DDR genes at low levels and contained mostly CN-low and grade 1 (G1) and grade 2 (G2) tumors (Fig. 1F). Tumors in Cluster 3 had DDR gene expression patterns strongly resembling those of ESC (Fig. 1F). Of the 37 DDR genes of TransSigGroup5 that characterized ESC, all were enriched in Cluster 3 endometrioid tumors as validated by Wilcoxon tests. Similar to ESC, high expression of TransSigGroup5 DDR genes in Cluster 3 endometrioid tumors was not associated with specific oncogenes or tumor suppressors. Also similar to ESC, Cluster 3 endometrioid tumors showed increased enrichment for TransSigGroup5 and 2 DDR genes compared to normal endometrium.

### Distinct endometrial cancer subsets based on DDR gene expression profiles are not driven solely by tumor grade effects

We considered the possibility that the high expression of TransSigGroup5 and 2 DDR genes associated with Cluster 3 endometrioid tumors was driven by tumor grade. However, when we removed the grade effect for each gene using linear models and re-analyzed DDR gene expression patterns, we still detected 3 main Clusters of endometrioid tumors (Supplementary Fig. 2). TransSigGroup5 and 2 DDR genes remained enriched in Cluster 3 tumors when tumor grade effect was removed (Fig. 1G). Approximately, 78% of Cluster 3 endometrioid tumors identified in Fig. 1F (without removal of tumor grade effect) remained in Cluster 3 after we removed the tumor grade effect (Supplemental Fig. 2). Similarly, most of the tumors remained in Clusters 1 and 2 after removing the tumor grade effect (Fig. 1G). Therefore, tumor grade alone cannot explain the separation of endometrioid tumors into 3 distinct Clusters based on DDR gene expression profiles. Expression of TransSigGroup5 and 2 DDR genes did not correlate with any particular oncogene or tumor suppressor mutations in EEC. HRD scores were not significantly different between the 3 endometrioid tumor clusters. However, Cluster 3 tumors showed higher RS and CA20 scores compared to Cluster 1 and Cluster 2 (RS score: Pearson’s chi-square statistic X is 4.49 for 1 degree of freedom, and p-value = 0.03; CA20 score: Fig. 1H). Therefore, TransSigGroup5 and 2 DDR genes enriched in endometrioid tumors (Cluster 3) might allow tolerance of genotoxicities resulting from DNA replication stress and supernumerary centrosomes.

### DDR gene expression profiles reveal distinct endometrial cancer subsets unrelated to molecular subtypes

Next, we asked whether TransSigGroup5 and 2 DDR genes showed any relationship with the endometrial cancer molecular subtypes identified by proteogenomic characterization.^7^ Therefore, we performed supervised clustering of both ESC and EEC based on the four TCGA genomic subtypes (namely CN-high, CN-low, MSI, and POLE-mutated) instead of histological subtypes. As shown in Supplemental Fig. 3, TransSigGroup5 DDR genes corresponded to the CNV-high subtype (a category which contains nearly all ESC). A subset of the POLE-mutated tumors and a subset of MSI tumors (two categories which contain endometrioid tumors) also were enriched for TransSigGroup5 DDR genes (Supplemental Fig. 3). Of the various classifiers we used, high tumor grade and CA20 scores were most closely associated with expression of TransSigGroup5 DDR genes in CNV-high, MSI, and POLE. The estimated Spearman Rho for CA20 score is 0.38 with a p-value < 0.001. Thus, genomic subtypes, even those related to DNA damage (MSI and POLE-mutated), cannot explain the DDR gene expression profiles of endometrioid tumors. We conclude that DDR gene expression profiles reveal distinct endometrial cancer subsets.

### ESC and Cluster 3 endometrioid tumors are defined by multiple DNA repair pathways and high CA20 scores

Most DNA repair processes rely on the coordinated actions of multiple DNA repair proteins that function in a common pathway. To complement our analysis of individual DNA repair genes in ESC and EEC, we performed pathway-based analyses to determine the overall status of different dedicated DNA repair processes. As shown in Fig. 2A our pathway-based analysis of DDR gene expression also revealed marked differences between ESC, EEC Clusters 1, 2, and 3, and normal endometrial tissues. In ESC and Cluster 3 EEC, we observed increases in expression of key DNA repair genes corresponding to multiple pathways. Thus, TMEJ, checkpoint signaling, FA, HR, BER and MMR pathways were generally elevated in ESC and Cluster 3 EEC when compared with normal endometrium or Clusters 1 and 2 EEC (Fig. 2A).

**Fig. 2.**
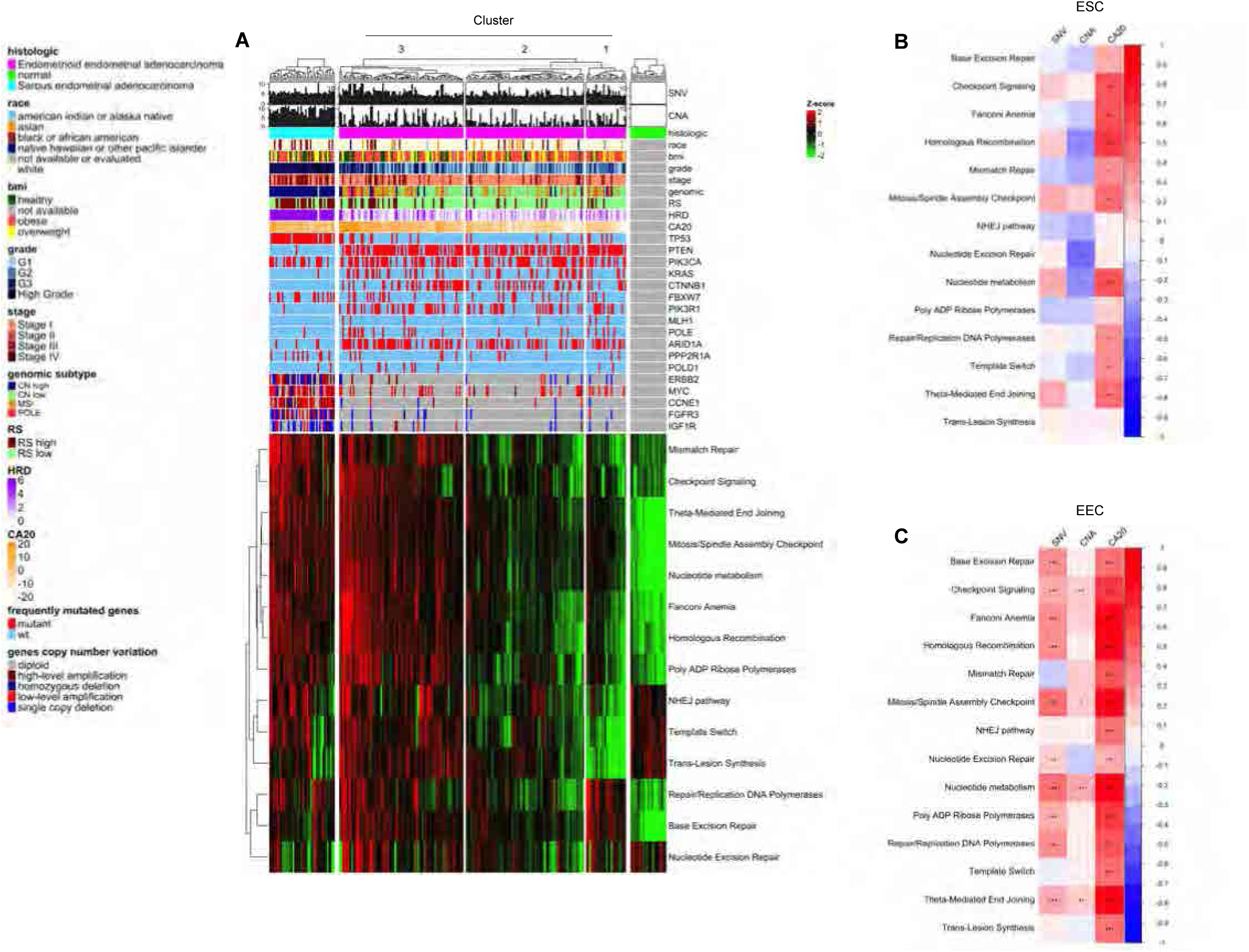
Comparison of serous and endometrioid endometrial tumors based on DDR pathway level analysis and SNV and CNA aberrations. (A) Supervised clustering of DDR pathway Z-scores by histology. K-means clustering of endometrioid tumors described in Fig. 2 was applied to endometrioid tumors after removing tumor grade effect. Spearman correlation matrix comparing DDR pathway Z-scores to CA20 score, total SNV counts, or CNV counts in (B) ESC and (C) EEC. (C-B) Spearman’s rank correlation. *p < 0.05, **p < 0.01, ***p < 0.001.

Many forms of DNA damage can be repaired by error-free or error-prone pathways. Therefore, DNA repair pathway choice and availability can impact genome stability. We tested correlations between DDR gene expression patterns at the pathway level and two specific forms of genomic instability: copy number alterations (CNA) and single-nucleotide variants (SNV). As shown in Fig. 2B and 2C, distinct DDR pathways were associated with SNV and CNA in ESC and EEC. For example, EEC with higher CNA generally had higher expression of DDR genes involved in TMEJ, nucleotide metabolism, and checkpoint signaling. SNV-high EEC showed enrichment of DDR genes that mediate BER, checkpoint signaling, the FA pathway, HR, mitosis/spindle assembly checkpoint regulators, NER, nucleotide metabolism, poly ADP ribose polymerases, and TMEJ. Conversely, ESC with CNA-high status showed reduced expression of genes involved in HR, NER, and nucleotide metabolism. We observed no correlations between DDR pathway gene expression and SNV-high ESC. For ESC and EEC, CA20 scores were positively correlated with expression of genes corresponding to multiple DDR pathways (Fig. 2B-2C). Notably, none of the 37 TransSigGroup5 and 20 TransSigGroup2 DDR genes overlap with CA20 signature genes. Thus, ESC and Cluster 3 endometrioid tumors may have a high dependency on these DDR pathways.

### TransSigGroup5 and 2 DDR gene signatures predict poor patient outcomes in endometrioid endometrial carcinomas

We asked whether our TransSigGroup5 or 2 DDR gene signatures were predictive of patient survival and/or disease recurrence. Because ESC and EEC are clinically different uterine tumors, we assessed overall survival and progression-free survival for each histological subtype. For ESC, high expression of TransSigGroup5 and 2 DDR genes were not predictive of overall survival or progression-free survival (Fig. 3A and 3C). These data are fully expected because ESC tumors are homogenous and all express similarly high levels of TransSigGroup5 and 2 DDR genes (Fig. 1A). In contrast with ESC, EEC are heterogenous with respect to expression of TransSigGroup5 and 2 DDR genes (Fig. 1A, Supplementary Fig. 1). Given that TransSigGroup5 and 2 DDR genes were enriched in approximately half of *POLE*-mutant EEC (Supplemental Fig. 3), *POLE*-mutants were removed in our Kaplan-Meier analyses for EEC. *POLE* mutations occur ∼9% of endometrial tumors and have excellent clinical outcomes even when tumors have high-risk features (e.g., high grade, lymphovascular space invasion, are an aggressive non-EEC histotype).^25,26^ Recent evidence suggests that *POLE* mutations override the biological relevance of other genetic/molecular alterations (e.g., p53) of the tumors.^27,28^ Interestingly, high expression of TransSigGroup5 and 2 DDR gene signatures in *POLE*- proficient EEC were predictive of poor patient outcomes for EEC. High expression of TransSigGroup5 or 2 DDR genes correlated with worse overall survival and progression-free survival of EEC patients (Fig. 3B and 3D). Expression of several individual genes from TransSigGroup5 and 2 was strongly associated with poor overall survival (18 genes) and/or worse progression-free survival (15 genes) (Supplemental Figs. 4 and 5). These results demonstrate that Transcriptome signatures Group 5 and 2 identify aggressive endometrial cancers, regardless of being CN high, CN low, or MSI subtypes.

**Fig. 3.**
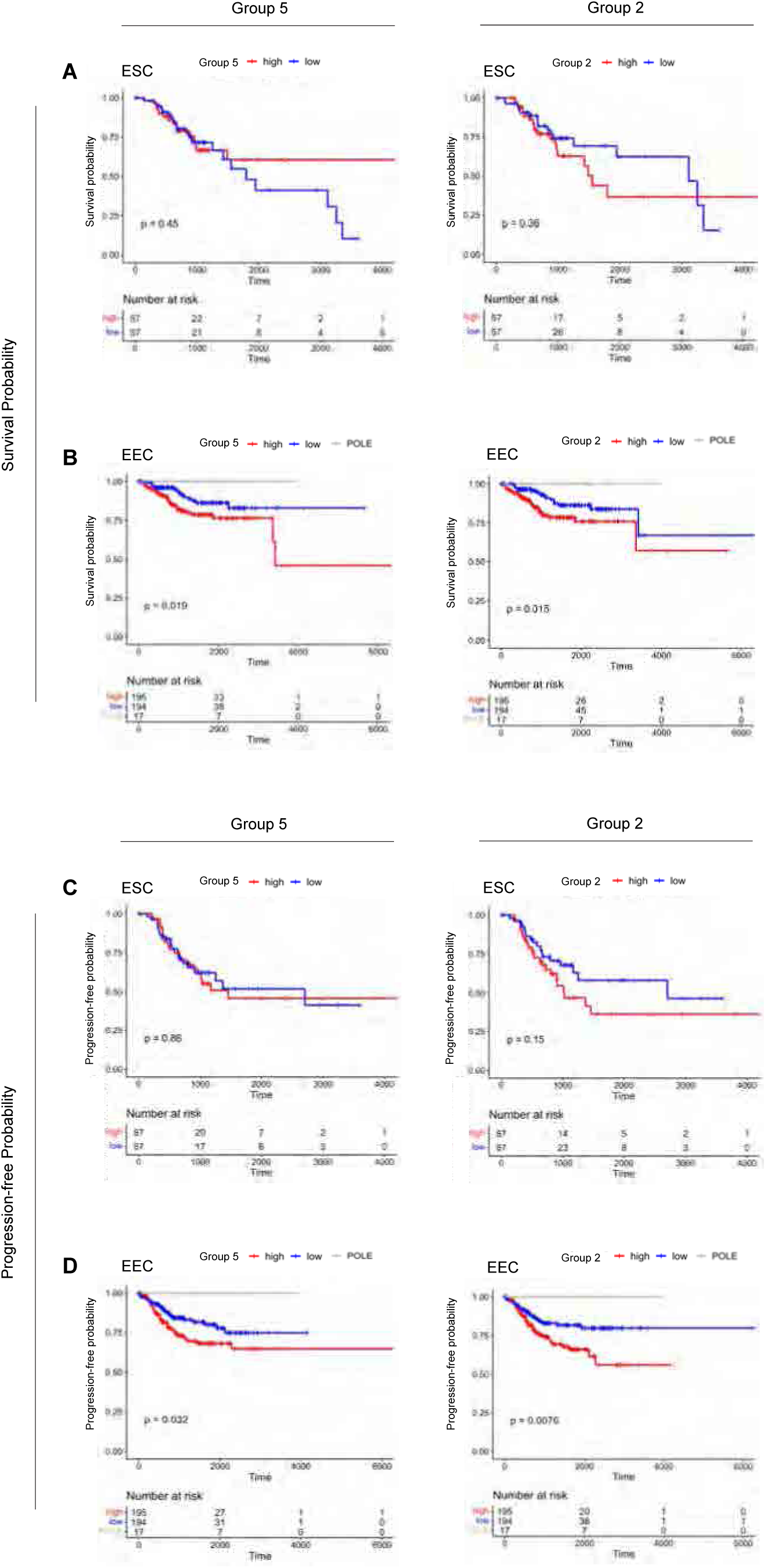
TransSigGroup 5 and 2 predict poor outcomes in endometrioid endometrial carcinomas. Kaplan-Meier curves for overall survival (A-B) and progression free survival (C-D) of TransSig(Group 5) and TransSig(Group 2) in ESC or EEC. Patient data used are from the entire uterine cancer cohort from the TCGA (similar to Fig. 2B). Patient cases ESC and non-*POLE* mutant EEC were divided into two groups (‘high’ and ‘low’) based on median TransSig(Group 5) or TransSig(Group 2) DDR gene expression. Log-rank tests were used to infer statistical significance between high and low groups.

### Endometrial cancer-defined DDR gene signatures and POLQ association are not generalizable to all HRD-high tumors

HR pathway deficiency is associated with several cancer types, including high grade ovarian serous carcinoma (HGOSC) and pancreatic adenocarcinoma (PDAC). Fig. 1A shows that HRD scores are uniformly high in ESC compared to low grade EEC. Based on histologic and genetic/molecular similarities and clinical behavior ESC is considered to be clinically equivalent to HGOSC^6,29^. Therefore, we determined whether TransSigGroup5 and 2 DDR gene signatures associated with ESC were also hallmarks of HGOSC and other HRD-high tumors (such as PDAC). Interestingly, ESC and a large subset of HGOSC shared similar enrichment Z-scores for TransSigGroup5 DDR genes (Supplementary Fig. 6A) and several DNA repair pathways (Supplementary Fig. 6B). In contrast, TransSigGroup5 DDR genes were expressed at low levels in PDAC (Supplementary Fig. 6A). Furthermore, expression patterns of all DDR genes in PDAC were distinct from those in ESC and HGOSC (Supplementary Fig. 6B). We conclude that expression of TransSigGroup5 and 2 DDR genes is not a byproduct of HR-deficiency, but instead is a specific characteristic of ESC and HGOSC.

Reportedly, in a subset of HGOSC with HRD features there is compensatory upregulation of DNA Polymerase theta (POLQ), the rate-limiting enzyme in the Theta-Mediated End-Joining (TMEJ) DSB repair pathway.^30^ Because POLQ is error-prone, TMEJ generates a microhomology deletion (MHD) mutational scar that is represented in COSMIC mutational signature 3.^31^ Given the clinical resemblance of ESC and HGOSC, we investigated links between HRD status, *POLQ* expression, and mutation signature 3 in endometrial cancer. We used the method of Marquard et al.^32^ to assess HRD status in high and low grade EEC, ESC, HGOSC, and PDAC. As shown in Supplementary Fig. 6C, ESC, HGOSC, and PDAC had relatively high HRD scores when compared with high and low grade EEC. In contrast with previous reports, we did not observe a correlation between *POLQ* expression and HRD status in HGOSC (Supplementary Fig. 6D). Similarly, *POLQ* expression did not correlate with HRD status in ESC (Supplementary Fig. 6D). We only observed positive associations between *POLQ* expression and HRD in EEC and PDAC (Supplementary Fig. 6D). We conclude that the expression of POLQ previously described in HGOSC^30^ cannot be attributed solely to HR-deficiency. COSMIC mutation signature 3 (whose etiology is attributed to HR-deficiency and may involve *POLQ*) was present in 45.2% (19/42) of ESC and 87.2% (198/227) of HGOSC. We found no correlation between COSMIC mutation signature 3 and *POLQ* expression with ESC or HGOSC (Supplementary Fig. 6E). Cosmic signature 3 was not detectable in most EEC or PDAC (data not shown) and therefore we did not test correlations with *POLQ* expression in those tumor types. Taken together, we show that the relationship between HRD status, POLQ expression and COSMIC signature 3 is more similar in clinically related gynecological cancers (ESC and HGOSC) than in the clinically distinct PDAC and EEC.

### CRISPR screening defines DDR dependencies of endometrial cancer

Our results suggest that profiles of DDR gene expression in endometrial cancer reflect the genome maintenance requirements for tolerating intrinsic genotoxic stresses of tumorigenesis. To formally identify the DDR genes pathways that sustain high grade endometrial cancer cells, we devised an unbiased genetic screening approach. We engineered a lentiviral vector for co-expressing sgRNAs and a Shield-1-inducible CAS9 (DD-CAS9). The resulting vector was used to construct a library of sgRNAs targeting 504 DDR genes (10 unique sgRNAs per gene) spanning every major DNA repair pathway and many stress response pathways. Using our library, we performed CRISPR dropout screens to identify DDR gene dependencies of HEC-50 (also known as HEC-50B) endometrial cancer cells. HEC-50 cells were originally derived from ascites of a recurrent stage 3, high grade endometrioid tumor^33^. In mice, HEC-50 cells form tumors with serous-like histology. Thus, HEC-50 cells model both ESC and high grade EEC. The design of our CRISPR screen is illustrated in Fig. 4A. Our CRISPR screen identified 28 DDR genes that were necessary for the growth of HEC-50 cells (Table 2). Based on stringent thresholds (log2FC) for sgRNA dropout, HEC-50 cells showed greatest dependency on *PRKDC, ATM, USP7, FANCA, RNF168, RAD51C, and MRE11A* genes (Fig. 4B). We selected some of these genes for independent validation experiments. The clonogenic survival assays in Fig. 4C confirm that *PRKDC*, *USP7*, *RNF168*, and *MRE11* are required for HEC-50 cell growth thereby validating DDR gene dependencies identified by our CRISPR screen.

**Fig. 4.**
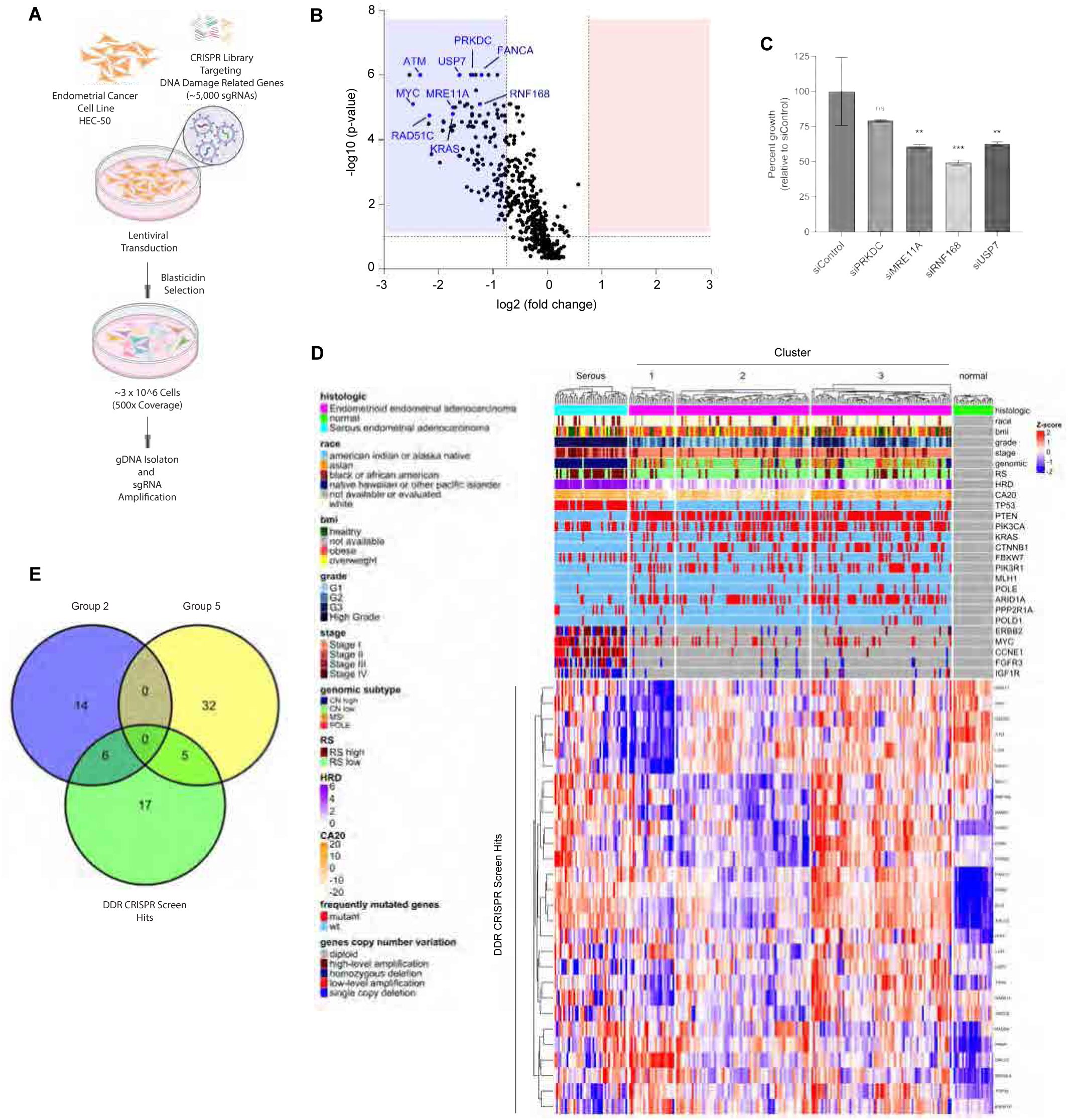
DDR CRISPR screen in high grade endometrial cancer cells reveals dependencies identified with TCGA transcriptome signatures. (A) Schematic illustration of CRISPR-Cas9 screen targeting DDR genes in HEC-50 cells. (B) Fold depletion in sgRNA targeting DDR genes analyzed by VOLUNDR pipeline. (C) Percent growth in HEC-50 cells after siRNA-mediated knockdown of indicated target genes, analyzed using clonogenic survival assay. (D) Expression patterns in patient tumors for the 28 DDR genes that dropped out (‘hits’) with CRISPR screens in HEC-50 cells. (E) Venn diagrams depicting DDR CRISPR screen ‘hits’ in HEC-50 cells compared to DDR genes in TransSig(Group 5) and TransSig(Group 2). (C) One-way ANOVA, Dunnett post-test, mean ± SD. *p < 0.05, **p < 0.005, ***p < 0.0005, ns = non-significant.

Next, we examined expression patterns in patient tumors and associations with clinical outcomes for the 28 DDR genes identified by our screen as HEC-50 dependencies (Fig. 6D). As shown in Fig. 4E, five of the DDR genes that were required for HEC-50 growth overlapped with the Group 5 gene list from Fig. 1A (*BLM, RRM2, FANCA, TIPIN, XRCC2*). Six of the DDR genes required for HEC-50 growth were present in Group 2 (*RAD51C, PRKDC, RRM1, RNF168, MDC1, BARD1*). Notably, half of these genes (*BARD1, PRKDC, RNF168, RRM2, FANCA*) correlate with low survival probability and/or worse progression-free survival in EEC patients (Supplemental Figs. 4 and 5), which is consistent with pro-tumorigenic roles for these DDR genes. Taken together, these results show that DDR genes expressed at high levels in ESC and a subset of EEC are associated with poor clinical outcomes and are necessary to sustain proliferation of endometrial cancer cells.

### ESC and Cluster 3 EEC share distinct CT Antigen expression patterns which are associated with genome maintenance and poor overall survival

Many CT Antigens (CTAs; germ cell proteins that are mis-expressed in cancers) pathologically activate DNA repair and confer DNA damage tolerance in tumors.^10,12,14^ The ways in which CTAs impact genome maintenance (and other tumorigenic characteristics) of endometrial cancers have not been defined. To determine potential roles of CTAs in genome maintenance and the pathobiology of endometrial cancer, we curated a list of 219 genes encoding known CTAs (Supplemental Table 2) and determined their expression patterns in endometrial tumors. The heatmap in Fig. 5A shows CTA expression patterns in relation to ESC and EEC clusters 1, 2, and 3 (as defined by DDR gene expression K means clustering in Fig. 1F). Unexpectedly, several CTAs were expressed in normal endometrial tissues, indicating that some genes previously designated as CTAs may also have roles in non-malignant tissues.

Our supervised clustering revealed a distinct set of 16 CTA genes that had very low expression in normal endometrial tissue but were highly expressed in ESC and EEC cluster 3 (Fig. 5A; Group 1). Those ‘Group 1 CTA genes’ are listed in Table 3. The elevated expression of Croup 1 CTA genes in EEC cluster 3 remained when tumor grade effect was removed (Supplemental Fig. 7). Group 1 CTA genes significantly correlated with HRD and CA20 scores (Figs. 5B-5C) and high RS signature (Fig. 5D), potentially consistent with genome maintenance-related roles for these CTA.

**Fig. 5.**
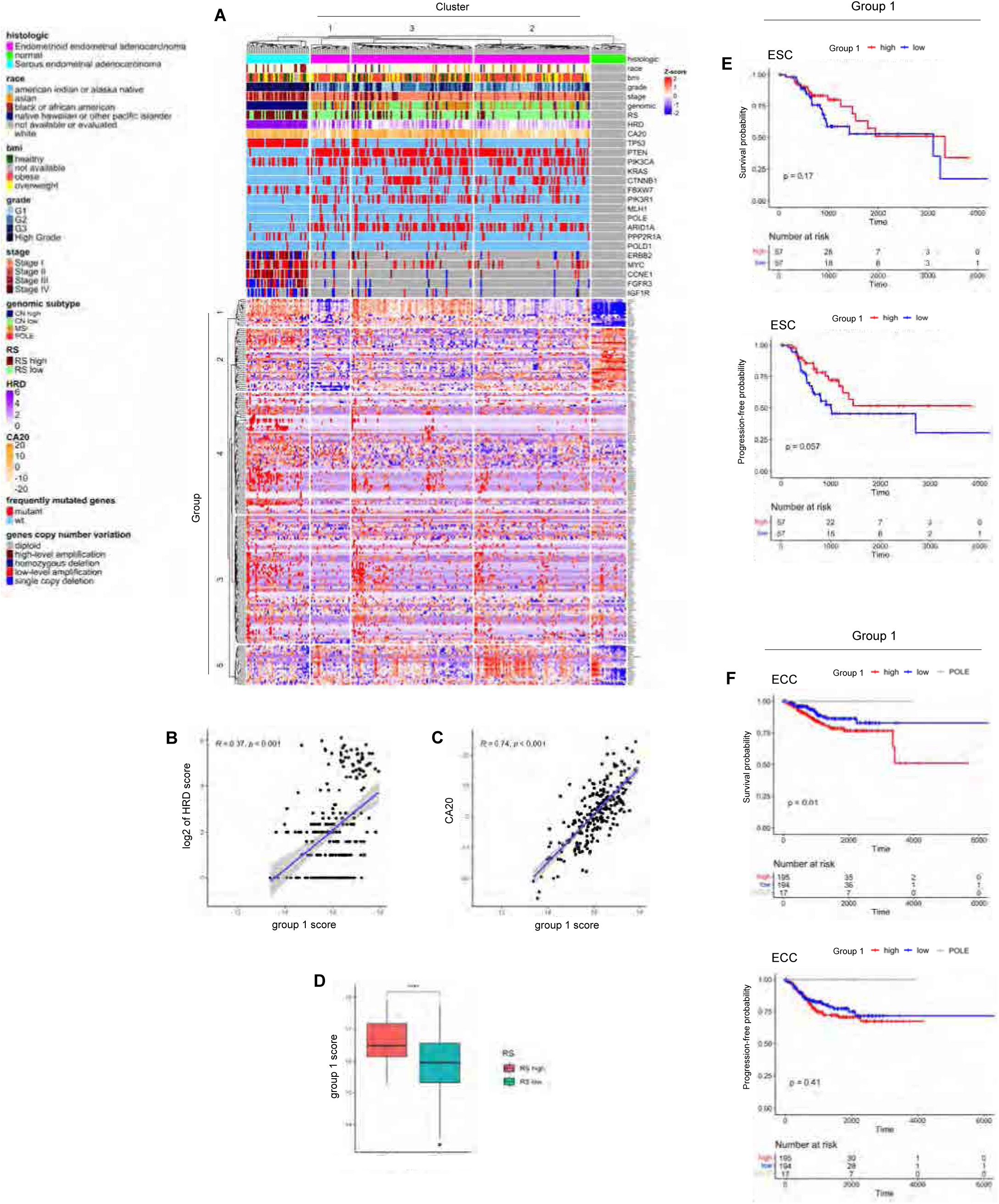
Expression of CT Antigen genes in endometrial cancer with supervised clustering based on histological subtypes. (A) Samples from the 2013 TCGA sample set for uterine cancer clustered by endometrial tumor histology (serous or endometrioid), normal endometrium, and K-means clustering of endometrial endometrioid carcinomas (defined in Fig. 2A). Heatmaps are depicting relative mRNA expression of CTA genes assembled from a curated list of 248 genes. 219 genes are displayed in the heatmap as 29 genes had no expression in normal endometrium or endometrial tumors. Transcriptome gene signature Groups 1-6, identified by K-means clustering of DDR gene expression in Fig. 1A, are depicted on the heatmap. Spearman’s rank correlation of CTA genes of Group 1 of all endometrial tumors vs. HRD (B) or CA20 (C) scores or by RS high or low status (D). Group 1 gene score is the average expression of the genes in the grouping. Kaplan-Meier curves for overall survival and progression free survival in ESC (E) and non-*POLE* mutant EEC (F). Patient data is from the entire uterine cancer cohort from the TCGA. Data were divided into two groups (‘high’ and ‘low’) based on median Group 1 CTA gene expression. Log-rank tests were used to infer statistical significance between high and low groups.

Group 1 CTA gene expression did not correlate with any particular oncogene or tumor suppressor mutations (Fig. 5A and Supplemental Fig. 7) or molecular subtype (Supplemental Fig. 8A). Similar to the TransSigGroup5 and 2 DDR gene signatures, high expression of Group 1 CTA genes was collectively predictive of poor overall survival in EEC and not ESC (Fig. 5E- 5F). Therefore, expression of these CTAs defines distinct disease subsets that cannot be captured by tumor grading, molecular subtyping, or histopathology. These data underscore the potential contribution of CTAs to tumorigenic properties of endometrial tumors.

Given the correlation of Group 1 CTA genes with high HRD scores, we asked whether these CTAs share a similar expression pattern in ESC and HGOSC (two clinically similar HRD-high tumors) and PDAC (which are HRD-high yet clinically dissimilar to ESC). Supplemental Fig. 8B shows that Group 1 CTA genes were expressed at similar high levels in ESC and HGOSC, but were present at very low levels in PDAC. Therefore, although expression of Group 1 CTA genes is associated with HR-deficiency in endometrial cancers, HRD status alone is insufficient to drive expression of Group 1 CTA genes in all cancers.

### Group 1 CTAs promote tumorigenic phenotypes in endometrial cancer cell lines

We tested the contribution of Group 1 CTA genes to tumorigenic behavior of endometrial cancer cells. First, we confirmed that several of the Group 1 CTA genes (*KIF2C, NUF2, CEP55*) were expressed at high levels in endometrial cancer cell lines when compared with normal endometrial epithelial cells (iHEuEC) (Fig. 6A). Knockdown of *KIF2C, CEP55,* and *NUF2* using siRNA reduced clonogenic survival of endometrial cancer cells (Fig. 6B), thus implicating a role for these CTAs in promoting a key tumorigenic characteristic. To test the relationship between CTAs and oncogene signaling we conditionally expressed endometrial cancer-relevant oncogenes (*CCNE1*, *KRAS^G12V^*, *MYC*) in normal endometrial epithelial cells (iHEuEC). As shown in Fig. 6C, induced expression of *CCNE1* and *KRAS* oncogenes led to increased expression of KIF2C, CEP55, and NUF2. Moreover, clonogenic survival of KRAS^G12V^-expressing cells (but not parental cells lacking KRAS^G12V^) was attenuated by treatment with NUF2 siRNA (Fig 8D). We conclude that CTA expression can be an adaptive response that confers tolerance of oncogene-associated mitogenic stresses.

**Fig. 6.**
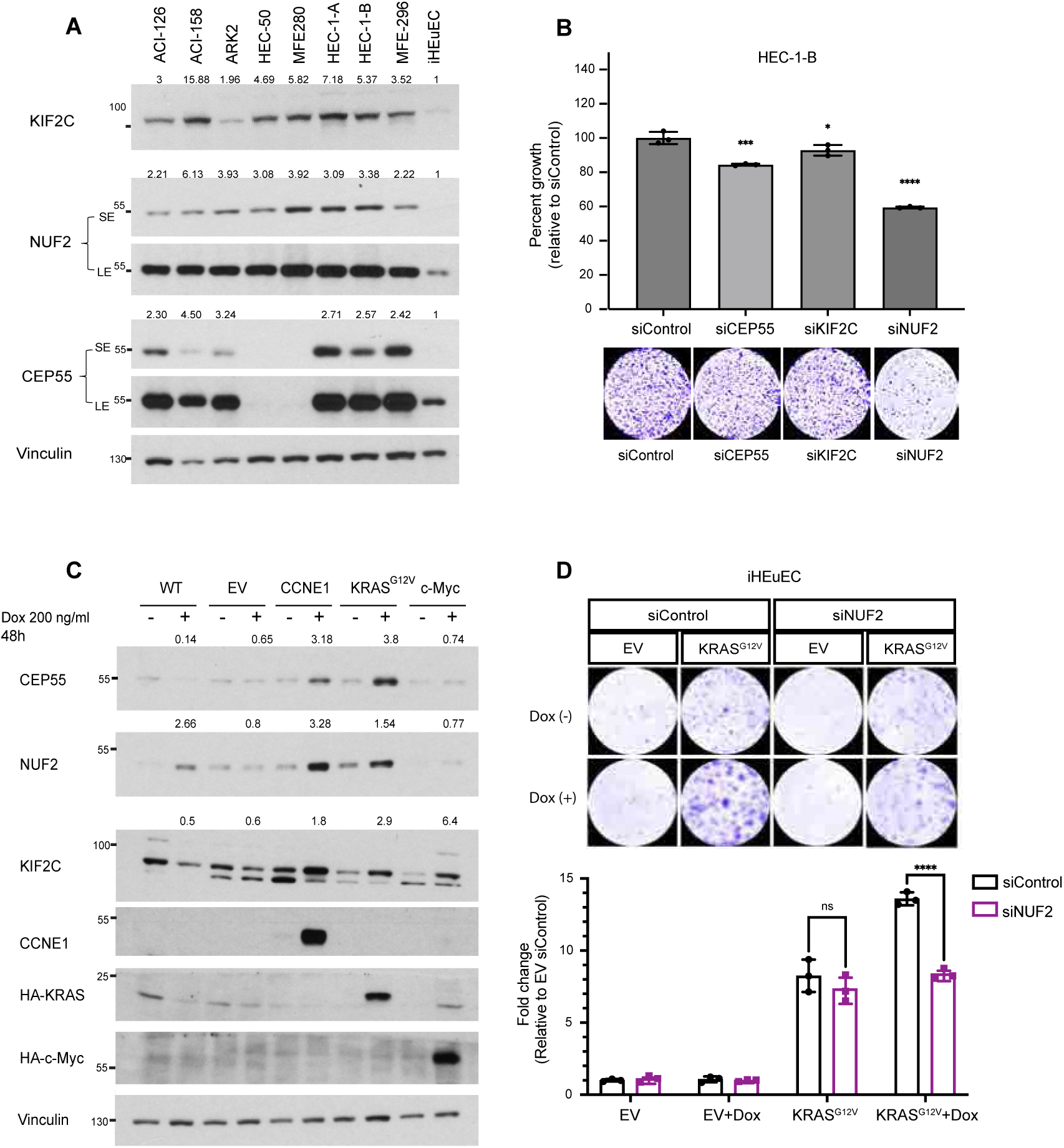
Endometrial cancer cell growth is dependent on Group 1 CTAs. (A) Immunoblots showing protein expression of CTAs KIF2C, NUF2, and CEP55 in immortalized human endometrial epithelial cells (iHEuEC) and various endometrial cancer cell lines. Numerical values in each lane indicate fold change in expression of indicated protein relative to its expression in iHEuEC cells. (B) Relative growth of HEC-1-B cells after siRNA-mediated knockdown of indicated CTA genes (KIF2C, CEP55 and NUF2) by clonogenic survival assay, quantified relative to control siRNA transfected cells. (C) Immunoblots showing protein levels of KIF2C, NUF2 and CEP55 in iHEuEC cells stably expressing Dox-inducible oncogenes *CCNE1*, *KRAS^G12V^* and *c-Myc*, relative to WT and iHEuEC empty vector (EV) transformed cells. iHEuEC cells were treated with Doxycycline (Dox) 200 ng/ml for 48h. Numerical values in each lane indicate fold change in expression relative to protein levels in their respective no Dox controls. (D) Relative growth of iHEuEC EV and iHEuEC expressing mutant *KRAS^G12V^* after transfection with either non-targeting Control (siControl) or siRNA targeting NUF2 (siNUF2). Relative growths were measured with or without Dox treatment (200 ng/ml) by clonogenic survival assay. Quantification of growth is relative to iHEuEC EV siControl cells (-Dox). (B) One-way ANOVA, Dunnett post-test, mean ± SD. (C) Two-way ANOVA, Dunnett post-test, mean ± SD. *p < 0.05, ***p < 0.0005, ****p < 0.00005.

### KIF2C allows endometrial cancer cells to tolerate DNA replication-stress

One of the most highly expressed Group 1 CTAs was KIF2C. This CTA encodes a kinesin-like motor that regulates the dynamic association of microtubules with kinetochores.^34^ The significance of high-level KIF2C expression in neoplastic cells, and the potential contribution of KIF2C to tumorigenic and/or chemoresistant phenotypes have not been addressed (for any malignancy). We asked whether KIF2C affects the responses of endometrial cancer cells to DNA-damaging events. In cells treated with Hydroxyurea (HU, an agent which depletes nucleotide pools and causes DNA replication stalling), KIF2C knockdown led to increased accumulation of pRPA and γH2AX (Fig. 7A). However, in cells treated with neocarzinostatin (NCS) or ultraviolet (UV) radiation (agents which induce DNA DSB and bulky DNA photo-adducts respectively) KIF2C-depletion did not cause persistent pRPA and γH2AX (Fig. 7A).

**Fig. 7.**
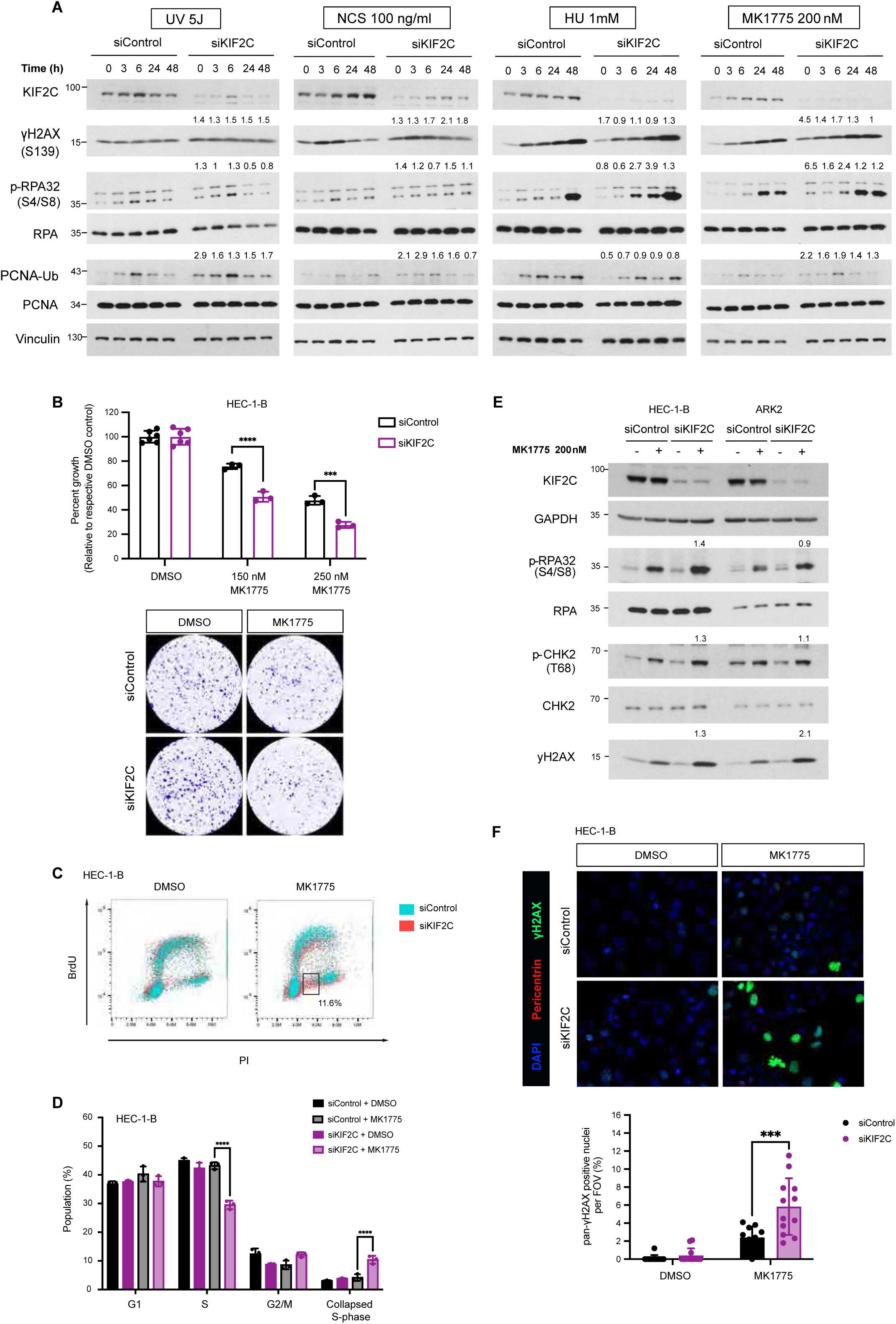
Loss of KIF2C sensitizes endometrial cancer cells to Wee-1-inhibiton-induced replication stress. (A) Immunoblots showing expression of DNA damage markers in crude chromatin lysates from HEC-1-B cells transfected with siControl or siKIF2C, treated with indicated genotoxic agents at different timepoints. Numerical values in siKIF2C lanes indicate fold change in expression, relative to respective siControl treatment group. (B) Relative growth of HEC-1-B cells after transfection with either non-targeting Control (siControl) or siRNA-targeting KIF2C (siKIF2C). Cells were treated with either DMSO or MK1775 (150 nM) for 72 hours. Growth was analyzed by a clonogenic survival assay. (C) Overlay of BrdU-PI cell cycle profile analyzed by flow cytometry of HEC-1-B cells. Cells were transfected with either siControl or siKIF2C, treated with either DMSO or MK1775 (200 nM) for 48 hours. (D) Quantification of percentage of cells in different phases of cell cycle (G1, S, and G2/M) along with S-phase defects that can be visualized as collapsed <4n populations. (E) Immunoblots showing expression of DNA damage markers in whole cell lysates from HEC-1-B and ARK2 cell lines transfected with siControl or siKIF2C, treated with DMSO or MK1775 (200 nM) for 48h. Numerical values in siKIF2C lanes indicate fold change in expression, relative to expression in respective siControl treatment group (after normalization to loading control or respective total protein). (F) Immunofluorescence images of HEC-1-B cells transfected with siControl or siKIF2C +/- MK1775 (200 nM) treatment for 48h. Transfected cells were synchronized with single thymidine block (2 mM) treatment. After thymidine release, cells are treated with MK1775 for 48h and then fixed for detection of immunodetection of γH2AX, pericentrin and DAPI. Quantification of pan-γH2AX positive nuclei in immunofluorescent images indicative of replicative stress in indicated groups. (B, D, and F) Two-way ANOVA, Tukey post-test, mean ± SD. ***p < 0.0005, ****p < 0.00005.

We also asked whether KIF2C protects against therapy-induced DNA damage. Inhibitors of the WEE1 protein kinase have been used as experimental therapeutic agents in patients with ESC.^35,36^ Mechanistically, WEE1 inhibition can lead to DNA replication stress, bypass of G2 checkpoints, and mitotic catastrophe.^18^ In HEC-1-B cells, KIF2C was recruited to chromatin after WEE1 inhibitor-treatment (MK1775; Fig 7A). MK1775-treatment led to more rapid and high-level accumulation of DNA damage markers pRPA and γH2AX in KIF2C-ablated HEC-1- B cells when compared with control (KIF2C-replete) cells (Fig. 7A).

HEC-1-B cells were also sensitized to MK1775-treatment following KIF2C depletion (Fig. 7B; Supplementary Fig. 9). In flow cytometry experiments, cells lacking KIF2C failed to maintain normal rates of ongoing DNA synthesis in the presence of WEE1 inhibitor (Fig. 7C and 7D; Supplemental Fig. 9F and 9G). In immunoblotting experiments, the increased levels of pCHK2, pRPA, and γH2AX expression in KIF2C-depleted MK1755-treated HEC-1-B and ARK2 cells are indicative of a role of KIF2C in remediating DNA replication stress (Fig. 7E). Immunofluorescence microscopy revealed pan-nuclear γH2AX in KIF2C-depleted and WEE1 inhibitor-treated cultures (Fig. 7F). The γH2AX present at sites of DNA DSB repair typically shows a pattern that is focal, not pan-nuclear. Therefore, the γH2AX staining pattern shown in Fig. 7F is indicative of DNA replication fork collapse, fully consistent with the reduced DNA synthesis rates of KIF2C-depleted cells shown in Fig. 7C and 7D. Taken together, we conclude that KIF2C is important for tolerance of a subset of endogenously arising as well as pharmacologically induced species of DNA lesions that arise in endometrial cancer cells during S-phase. Moreover, our results indicate that KIF2C, CEP55, NUF2, and likely other CTA confer key tumorigenic properties and contribute to the pathobiology of endometrial cancers.

## Discussion

Here, we present the first comprehensive characterization of the DDR and CTA transcriptional landscape of endometrial cancer in relation to conventional histological and molecular subtypes. We show that endometrial tumors have distinctive DDR and CTA expression profiles when compared with other tumor types. Critically, DDR and CTA expression patterns of endometrial cancers associate strongly with clinical outcomes. In fact, DDR-based profiling is prognostic of endometrial cancer patient outcomes in ways that other classification methods fail to capture. Historically, it has been observed that a subset of EEC behave very similarly to ESC (clinical aggressiveness and poor outcomes), despite having different histological and molecular characteristics. Defining these aggressive EEC tumors has been challenging and even high tumor grade (grade 3) has not always been a reliable indicator of EEC patients with aggressive disease. Remarkably, our studies reveal that aggressive *POLE*-proficient endometrioid tumors with poor outcomes can be clearly defined (and shown to resemble the ESC subtype) based on DDR and CTA expression profiling. Thus, DDR and CTA expression-based classifiers (specifically TransSigGroup5 and TranSigGroup2) identify aggressive, endometrioid tumors very consistently and independently of histology and molecular subtype and these signatures are not driven solely by tumor grade effects.

Why then do DDR and CTA expression profiles represent such robust and reliable biomarkers for tumor classification? Neoplastic cells exist in various stressful environments and must tolerate unique genotoxic stresses, while acquiring specific genomic changes that propel tumorigenesis. The DDR allows cells to withstand genotoxic stresses. Moreover, deployment of error-prone DNA repair pathways can promote genomic instability. Our work suggests that the unique stresses experienced by different endometrial cancer subtypes may necessitate specific DDR dependencies that are accurately reflected by their DDR transcriptomes.

The two DDR mRNA gene signatures that are strongly associated with ESC and the aggressive EEC subtypes (TransSigGroup2 and 5, comprising 20 and 35 DDR genes respectively) have some interesting and noteworthy characteristics: TransSigGroup2 and 5 contain the *RRM1* and *RRM2* transcripts which encode subunits of Ribonucleotide Reductase (RNR), an enzyme essentially required for the conversion of rNTPs into dNTPs that sustain DNA synthesis and repair^37^.

TransSigGroup5 contains mRNAs encoding all three members of the Replication Fork Protection Complex (TIM, TIPIN, CLASPIN) which stabilizes DNA replication forks and promotes S-phase checkpoint signaling^38^. TransSigGroup2 and 5 also contain genes encoding the S-phase Checkpoint kinase CHEK1 and its activators TOPBP1 and RAD1^39^. Genes encoding other important mediators of DNA replication fork progression are also present in TransSigGroup2 and 5 including the Trans-Lesion Synthesis (TLS) factors RAD18 and POLH, and the Template Switch (TS) factor HLTF (which cooperates with RAD18)^40^. In TransSigGroup5 and 2 gene lists there is very strong representation of mRNAs encoding mediators of the Fanconi Anemia FA) and Homologous Recombination (HR) pathways (Table 1) which also facilitate replication fork progression in times of stress.

Our CRISPR screen validated many TransSigGroup2 and 5 DDR genes (and their associated pathways) as important dependencies of cultured endometrial cancer HEC50 cells. Genes identified by our CRISPR screen include, the RNR subunits RRM1 and RRM2, the RFPC component TIPIN, and RAD9A and HUS1 (whose encoded proteins associate with RAD1 to form the 9-1-1 complex that activates CHEK1). CRISPR screening also confirmed requirements for FA and HR pathway genes including BARD1, FANCA, XRCC2 (FANCU), RAD51C and RAD51D. RNF168 (identified by our CRISPR screen) is not a designated FA gene, but regulates localization of BARD1 which in turn recruits the BRCA1-PALB2 complex to DNA damage.^41^ Taken together, our DDR expression profiling and unbiased CRISPR screens suggest that high-grade endometrial cancer cells are particularly dependent on a subset of DDR factors involved in dNTP production, replication fork protection, the FA pathway and HR.

The high degree of overlap between the DDR genes overexpressed in TCGA samples and those identified by our CRISPR screen is remarkable considering that cancer cell growth in culture may create very different dependencies from those that arise in an *in vivo* pathological environment. Many of the FA pathway genes that were overexpressed in patient tumors were not required for growth in cell culture. In humans the FA pathway is particularly important for tolerance of acetaldehyde genotoxicity.^42^ We speculate that metabolic differences between tumors growing *in vivo* and cells in culture might explain the lack of complete overlap between TransSigGroup2 and 5 genes and dependencies identified by CRISPR screens.

Oncogene-induced DNA replication stress represents a major source of intrinsic DNA damage in cancer cells.^11,17,40,43–45^ Diverse mechanisms have been proposed to explain how oncogenes induce DNA replication stress (including re-replication, hyper-transcription and R-loop collisions, reactive oxygen species, and altered replication licensing). Moreover, different oncogenes may induce different species of DNA damage. Therefore, we predicted that DDR gene expression patterns might correlate with specific RS-inducing oncogenic drivers and/or with a composite ‘RS’ signature. Consistent with this prediction, the more aggressive ESC subtype (which expressed high levels of TransSigGroup2 and 5 DDR genes) did generally have stronger RS signatures when compared with all other endometrial patient tumors. However, in the aggressive Cluster 3 EEC subset (which similar to ESC expressed TransSigGroup2 and 5 genes), DDR gene expression patterns did not correlate with alterations in any of the *KRAS*, *MYC*, *CCNE1*, *PTEN*, *TP53*, *PIK3CA*, *FBXW7*, *CTTNB1* or with the RS signature. However, TransSigGroup5 and 2 DDR genes (expressed in ESC and aggressive EEC subsets) correlated strongly with the CA20 centrosome amplification signature.

Centrosome amplification occurs in many cancers and contributes to genomic instability and tumor initiation^22,46–49^. The formation of a normal bipolar mitotic spindle strictly requires the presence of two centrosomes at mitosis. Excessive numbers of centrosomes can cause spindle multipolarity, and mitotic delay leading to DNA damage through direct and indirect mechanisms^50–55^. Therefore, cancer cells with supernumerary centrosomes must adapt to withstand the stresses and DNA damage caused by multipolar spindles. Altered transcriptional programs such as upregulation of TransSigGroup2 and 5 DDR genes is a plausible explanation for how EEC tolerate DNA damage caused by supernumerary centrosomes in CA20-high tumors.

In both ESC and EEC, high CA20 scores also correlated strongly with expression of a subset of 10 CTA genes (which we designated ‘Group 1 CTAs’). Genes represented in the Group 1 CTA list are *CNOT9*, *ATAD2*, *PBK*, *KNL1*, *CEP55*, *KIF20B*, *TTK*, *NUF2C*, *KIF2C*, and *OIP5*. While two of the Group 1 CTA gene products have roles in gene expression (CNOT9 regulates mRNA degradation^56^ and ATAD2 is a transcriptional regulator^57^), most have roles in mitosis: NUF2 and OIP5 are centromeric proteins with roles in chromosome segregation^58,59^. PBK is a mitotic protein kinase that mediates DNA damage-induced G2/M checkpoint control^60^. TTK (MPS1) encodes a dual specificity protein kinase that is essential for chromosome alignment and segregation during mitosis^61^. CEP55 (Centrosomal Protein 55) regulates mitotic cytokinesis^62^. KNL1 is a TTK/MPS1 substrate required for creation of kinetochore-microtubule (MT) attachments and chromosome segregation^63^. KIF20B and KIF2C are MT motor enzymes required for cytokinesis^64^. Remarkably, we show that several Group 1 CTA are oncogene-inducible and required for growth of endometrial cancer cell lines. Taken together our results suggest that Group 1 CTA are very important for regulating chromosome segregation and remediating mitotic stresses experienced by endometrial cancer cells with supernumerary centrosomes. Interestingly, other CTAs have also been implicated as important requirements for mitotic progression of lung cancer cells^65^.

Because Group 1 CTA are absent from normal somatic cells, these proteins are attractive as therapeutic targets for sensitizing cancer cells to mitotic stress. Importantly, CTA-directed therapies would be innocuous to normal healthy cells (which lack CTA expression). Two of the Group 1 CTAs identified here (PBK and TTK) are protein kinases and are druggable^61^. The other CTAs in Group 1 have not been evaluated for chemical tractability. However, the MT motors KIF2C and KIF1B use ATP as a substrate and therefore might well be druggable based on ATP-binding site competitive inhibitors. Mitotic stress can result from intrinsic defects as well as microtubule poisons such as taxanes^66^. Microtubule inhibitors are commonly used in endometrial cancer chemotherapy regimens. The mechanism of anti-mitotic agents such as paclitaxel is attributed to induction of spindle multi-polarity.^66^ Therefore, therapies targeting Group 1 CTAs might also be useful for sensitizing endometrial cancers to paclitaxel and other microtubule inhibitors. Taken together, the DDR and CTA landscape revealed by our study provides new opportunities for therapies that target vulnerabilities of aggressive endometrial cancer subtypes. Our study fully validates the use of DDR and CTA-based tumor classifiers and justifies extension of this classification method to other cancer types.

## Methods

### Data availability

TCGA-UCEC, TCGA-OV and TCGA-PAAD datasets were downloaded from TCGA portal (https://portal.gdc.cancer.gov/). Integrated mRNA expression data as well as HRD score mRNA expression files (FPKM-UQ) can be found at Xena browser (https://xenabrowser.net/). The Xena platform was used for visualizing and analyzing TCGA genomics data^67^. In analyses using the TCGA-UCEC datasets, we excluded samples with mixed serous and endometrioid histology. In analyses with the 2013 TCGA-UCEC dataset^6^, we excluded samples without defined molecular subtypes (POLE, MSI, CN-high, CN-low). For TCGA-OV, our analyses only included high grade (G3 or G4) serous ovarian carcinoma.

### Heatmap Classifiers

Classifiers were derived through clinical information, somatic variant calling files, and copy number variation files using the GISTIC2 pipeline. Histology, race, BMI, stage, and tumor grade were obtained directly through clinical data available by the TCGA. Genomic subtypes and genes reported to be frequently mutated or having copy number variations in endometrial cancer were derived through available data from the 2013 TCGA study^6^. HRD score was derived as an unweighted sum of three signatures: Number of telomeric Allelic Imbalances (NtAI), Large-scale State Transitions (LST), and HRD-LOH^68^. Replication Stress (RS) was derived as previously described^69^. In brief, tumors classified as RS-high included at least one of the following alterations (*CCNE1* amplification, *MYC* and *MYCL1* amplification, *KRAS* amplification, *NF1* mutation, *RB1* two-copy loss, *CDKN2A* two-copy loss, *ERBB2* amplification) whereas tumors without these features were classified as RS-low. CA20 score was defined as the sum of the expression of 20 genes (log2 median centered). These genes include: *AURKA, CCNA2, CCND1, CCNE2, CDK1, CEP63, CEP152, E2F1, E2F2, LMO4, MDM2, MYCN, NDRG1, NEK2, PIN1, PLK1, PLK4, SASS6, STIL* and *TUBG1*^24^.

### Visualization and statistical analyses

R (version 4.1.0) was used for visualizing and data analyses. Heatmaps were generated using the R package ComplexHeatmap (Version: 2.8.0)^70^ function Heatmap() and separated by the EEC, ESC, and normal samples using the option column_split. The standardized log2 (fpkm + 1) was used as the expression value for genes in the heatmaps. The DDR pathway heatmaps shown in Fig. 2 were derived by calculating the average expression of each gene for the pathways. Functions, column_km() and row_km(), were used to perform k-means clustering for the DDR genes (indicated as Groups 1-6; Fig. 1A and 1F) and endometrioid tumors (indicated as Clusters 1-3; Fig. 1F and 2A). To remove tumor grade effect, univariate linear regression was applied to each gene with grade as the only covariate to calculate the coefficients of grade for each gene in endometrioid tumors. Serous tumors and normal endometrial tissue were removed when performing the linear model. Survival analyses were performed using the following R packages: survival (Version 3.2-13) and survminer (Version 0.4.9). Kaplan-Meier curves were constructed for visualizing the survival data and log rank tests used for hypothesis testing.

### Cell lines

HEC-1-B, HEC-50, and ARK2 cells were cultured in RPMI-1640 containing 10% fetal bovine serum (Gibco, #26140095) and 1% Penicillin-Streptomycin (Gibco, #15140122). Human Endometrial Epithelial Cells (HEuEC) cells were purchased from (Lifeline Cell Technology, #FC-0078) and cultured in DMEM:F12 medium containing 10% FBS, 1% Penicillin-Streptomycin, and 1% Insulin-transferrin-selenium (Gibco, #41400045). Doxycycline-induced oncogene expressing immortalized-HEuEC (iHEuEC) were cultured in DMEM:F12 media supplemented with 10% tetracycline negative FBS (GeminiBio, #100-800-500) and 1% Penicillin-Streptomycin. All cells were cultured at 37°C in humidified chambers with 5% CO2.

### Immortalization of Human Endometrial Epithelial Cells (HEuEC)

HEuEC were immortalized by overexpressing hTERT upon infection with retrovirus encoding hTERT cDNA. Retrovirus was generated from canine sarcoma D17 packaging cell line stably transduced with the MoMLV-pBABE-hygro-hTERT vector. Supernatants from this retroviral packaging cell line were collected when cells were 70-80% confluent. Supernatants was centrifuged at 500 x g for 5 min. and then filtered through 0.45 µm nylon filter (Corning, #431225). HEuEC cells were seeded in 6-well plates, infected at 50% confluency with 2 ml/well of hTERT-encoding retrovirus containing medium with 8 µg/ml polybrene, and allowed to incubate with the retrovirus for 48 hours. Successfully transduced cells were selected with 100 µg/ml Hygromycin B (Corning, #30240CR) for 14 days. Cells obtained were immortalized and labelled as iHEuEC cells.

### Plasmids and RNA interference

siRNA oligonucleotides were used for targeting specific genes transfected into cells using Lipofectamine 2000 as per manufacturer’s instructions. The following siRNAs were purchased from Horizon discovery: siKIF2C#1 (#L-004955-00-0005), siNUF2 (#L-006893-01-0005), siCEP55 (#L-005289-00-0005), siKIF2C#2 (#J-004955-06-0005), siKIF2C#3 (#J-004955-08- 0005), siRNF168 (#M-007152-03-0050), siMRE11A (#M-009271-01-0050), siUSP7 (#L-006097-00-0005), siPRKDC (#L-005030-00-0005), siControl (#D-001810-10-05). The lentivirus Doxycycline-inducible expression system using the pInducer20 vector was purchased from Addgene (#44012). pInducer20 constructs were cloned with *KRASG12V-HA* cDNA insert by HiFi Assembly (NEB, #E2621S). pInducer20-*CCNE1* construct was cloned using Gateway cloning (Invitrogen, #11791020). pInducer20-*cMyc-HA* vector was constructed by VectorBuilder (Vector ID: VB200317-7039asg).

### Generating doxycycline inducible oncogene expressing cells

Low-passage HEK 293T cells were seeded in 10 mm dishes to 60% confluency. At 24 hrs. post seeding, cells were transfected using Lipofectamine 2000 (Invitrogen, #11668019). Each dish was transfected with 6 µg lentiviral packaging plasmids (4.5 µg dNRF and 1.5 µg pMDK64) and 6 µg pInducer20-plasmid encoding oncogene cDNA (c-Myc-HA, KRAS-HA, CCNE1, or pInducer20 backbone). Plasmid DNA mix was diluted in 350 µl serum-free Opti-MEM media (Gibco, #31985070). In a separate reaction mix, 35 µl Lipofectamine 2000 was diluted in 350 µl of Opti-MEM media. Both DNA mix and lipofectamine solution were mixed gently and incubated for 15 mins. at room temperature. The DNA-lipofectamine mix was then added to cultured HEK293T cells which were incubated overnight. Approximately 16 hrs. post transfection, transfection medium was removed and replenished with 10 ml fresh medium. Lentivirus containing cell culture supernatant was collected at 24 hrs. and again at 48 hrs. Collected medium was centrifuged at 500 x g for 5 mins. to remove cellular debris and the supernatant was filtered through a 0.45 µm nylon syringe filter. Filtered lentivirus containing medium was stored at −80°C. To generate doxycycline-inducible oncogene expressing cell line iHEuEC, cells were seeded in 6-well plates. At 50-60% confluency, cells were incubated with 2 ml lentivirus-containing media mixed with 0.8 µg/ml polybrene for 24 hrs. After lentiviral infection, cells were selected with Geneticin (Gibco, #10131035) at 1000 µg/ml for 5 days. pInducer20 backbone without oncogene cDNA insert was used for generating empty vector controls.

### Immunoblots

Chromatin protein extracts were isolated from cultured cells using CSK lysis buffer (CSK buffer; 10 mM Pipes, pH 6.8, 100 mM NaCl, 300 mM sucrose, 3 mM MgCl2, 1 mM EGTA, 10 mM NaF, 25 mM β-glycerophosphate, and 0.1% Triton X-100) containing fresh protease inhibitor cocktail (Roche, #4693159001) and phosSTOP (Roche, #4906837001). Harvested cell lysates in CSK buffer were incubated on ice for 15 min., followed by centrifugation at 4000 rpm for 4 min. The supernatant was collected as soluble cell fraction and the pellet was washed with 1 ml CSK buffer at 4000 rpm for 4 min. The remaining pellet was re-suspended in minimal volume of CSK buffer which is sonicated, followed by nuclease treatment, and collected as crude chromatin fraction. For whole-cell protein extracts, cultured cells were washed with cold 1X PBS and cold CSK lysis buffer added to cultured monolayers. Cells were collected in lysis buffer by scrapping, lysates generated by sonication and then centrifugation at 20,000 x g for 15 mins, and supernatants collected. Proteins extracts were resolved on Tris-Glycine gels (Invitrogen, #XP04205BOX) by electrophoresis and transferred to a nitrocellulose membrane. Membranes were incubated for 1 hr. in 1X Tris-buffered saline containing 0.1% Tween 20 (TBS-T) and 5% non-fat milk (blocking buffer) and then incubated with primary antibodies (dilutions ranging from 1:500 – 1:2000), prepared in blocking buffer, at 4°C overnight. Membranes were washed with 1X TBS-T and then incubated with HRP-conjugated secondary antibody diluted in 1X TBS-T at 1:5000 for 1 hr. Chemiluminescent signals were captured using Perkin Elmer Western Lightning Plus ECL reagent and autoradiography film. Primary antibodies used included: γH2AX S139 (Millipore, #05-636); KIF2C (Abcam, #187652); phospho-RPA32-S4/S8 (Bethyl, #A300245A), RPA34 (Millipore Sigma, #NA18) CEP55 (Santa Cruz, #sc-374051); NUF2 (Abcam, #ab180945); HA-tag (Santa Cruz, #sc-7392); c-Myc (Santa Cruz, #sc-40); CCNE1 (sc-248); Vinculin (Sigma, #V9131; GAPDH (Santa Cruz, #32233); phospho-CHEK2-T68 (CST, #2661L); CHK2 (CST, #2662); PCNA (Santa Cruz, #sc-56).

### BrdU labeling and flow cytometry

Cultured cells were incubated with 10 mM BrdU for 1 hr. before trypsinization. Trypsinized cells were washed with 1X PBS and re-suspended in 65% DMEM with 35% ethanol for fixation overnight at 4°C. Fixed cells were denatured using HCl and then neutralized with 0.1 M Sodium Borate (pH 8.5). Cells were stained with a FITC conjugated anti-BrdU antibody for 1 hr. Following BrdU staining, cells were incubated in 1X PBS containing 10 µg/mL of propidium iodide (PI) and 8 µg/mL of RNaseA overnight at 4°C. Stained cells were analyzed for replication dynamics (BrdU staining) and cell cycle phase (PI staining) using an Accuri C6 plus flow cytometer (BD) and analyzed using the FlowJo software.

### Clonogenic growth assay

Cells were reverse transfected with siRNA using Lipofectamine 2000 overnight in 6 mm dishes. After transfection, the media on the cells was changed to fresh media. After 6 hrs., cells were seeded in 6-well plates at a seeding density of 2000 cells per well. Twenty-four hrs. after seeding, cells were either treated with MK1775 or DMSO for 6-7 days of clonogenic growth. Medium changes occurred every 72 hrs. After 6-7 days, culture plates were stained with 0.05% crystal violet in 1X PBS containing 1% methanol and 1% formaldehyde. Plates were scanned and colony area measurements calculated using ImageJv2 software.

### Immunofluorescence

HEC-1-B cells transfected with siControl or siKIF2C#3 were plated into 6-well plates with glass coverslips. Twenty-four hrs. after seeding, HEC-1-B cells were treated with thymidine 2 mM overnight and then released from the thymidine block into treatment with MK1775 200 nM for 48 hrs. The cells were then washed with 1X PBS and fixed using 4% PFA for 15 mins. at room temperature. Fixed cells were incubated in blocking solution (DPBS with 3% BSA and 5% donkey serum) for 1 hr. followed by incubation with primary antibodies (diluted 1:200 in blocking solution) overnight at 4 C. Cells were washed 3x with 1X DPBS for 5 mins. For detection of the primary antibodies, cells were incubated 1 hr. with secondary antibodies (Alexa Fluor 488 donkey, and Alexa Fluor 647 Goat anti-mouse, 1:400 dilution). Secondary antibodies were removed by washing 3x with 1X DPBS for 5 mins. and coverslips mounted to microscope slides using Prolong Gold mounting medium with DAPI (Invitrogen, #P36941). Antibodies: γH2A.X (Millipore, #05-636) and Pericentrin (Abcam, #ab4448).

### CRISPR-Cas9 based DNA Damage Repair (DDR) gene targeting library preparation

The pooled CRISPR-Cas9 library includes 5040 sgRNAs targeting 504 DNA Damage Repair genes (each gene has 10 independent targeting sgRNAs) and 1000 non-targeting control sgRNAs. The oligonucleotide pool encoding sgRNAs in the library was synthesized by CustomArray Inc. The DDR-CRISPR library was PCR amplified with Q5 high-fidelity polymerase. The amplified and PCR purified DDR-CRISPR library was cloned into LentiCRISPRv2-DD-Cas9-BSD by NEBuilder HiFi DNA assembly. Endura Electrocompetent cells (LGC Biosearch Technologies, #602422) were transformed with the DDR-CRISPR library by electroporation. Transformed bacteria were plated on twelve 15 cm LB ampicillin plates and incubated overnight at 37°C. LB media was added to each plate and the transformed bacterial colonies were recovered using sterile scrapers. Pooled bacterial colonies from 3 plates were transferred to 500 mL LB and grown with shaking at 37 °C for 3 hours. The DDR-CRISPR library plasmid DNA was purified from transformed bacterial culture using a Qiagen EndoFree plasmid DNA Maxi kit (Qiagen, #12362).

### DDR targeting CRISPR-Cas9 screen in HEC-50 cells

HEK293T cells were transfected with DDR-CRISPR-Cas9 pooled library plasmid using Lipofectamine 2000. Lentivirus containing media supernatant was harvested from transfected cells at 24 hrs. and 48 hrs. after transfection. Lentivirus containing media was centrifuged at 500 x g for 5 mins. and filtered through 0.45 µm nylon filters. Aliquots were frozen at -80C. HEC-50 cells were transduced with DDR-CRISPR-Cas9 library lentivirus at an MOI of ∼0.3. Lentivirus infected cells were treated with Blasticidin (10 µg/ml) for 4-5 days to select stably transduced cells. After selection, cells were treated with Shield-1 (1 µM) for 4 days. 3×10^6 cells were harvested at this timepoint (population doubling 0 (PD0)) for isolation of genomic DNA and for using as library controls after sgRNA enrichment. Another 3×10^6 cells were seeded in three 150 mm cell culture dishes for subsequent passages. Cells were passaged every 3-4 days. At the final passage (population doubling 20 (PD20)), 3×10^6 cells were harvested for genomic DNA isolation and sgRNA enrichment. Assuming a harvest of 6 µg genomic DNA content from 1×10^6 cells, library preparation was performed with 18 µg genomic DNA, thereby preserving ˜500 fold representation of each sgRNA. sgRNA were then PCR amplified in two steps: 1. sgRNA was PCR amplified using NEB Q5 Hot Start polymerase (NEB, #M0494L). The product was purified using AMPure XP beads (Beckman Coulter, Cat.no: A63881). 2. A PCR step was then used to add indexed sequencing adapters. Each secondary PCR reaction contained 1 ng of purified PCR product from the first PCR step as template in a 100 µL reaction volume containing indexing primers. After purification using AMPure XP beads, the 320 bp products were quantified using Qubit and diluted to 1 ng/mL and sequenced using HiSeq 4000, PE 2×150 (Novogene). Analysis of sgRNA enrichment was performed using Volundr pipeline.

## Supplemental Figure Legends

**Supplemental Fig. 1. Validation of expression patterns of TransSig (Group 5 and Group 2) DDR genes in a second cohort.** DDR gene expression patterns in all serous and endometrioid tumors in the TCGA database for uterine cancer (n = 521). Differential DDR gene expression patterns (Groups 1-6) identified in Fig. 1A are the same differential expression patterns seen with the larger cohort of endometrial tumors.

**Supplemental Fig. 2. Expression of DDR genes with removal of grade effect in endometrioid endometrial carcinomas.** Subtraction of tumor grade effect for endometrioid tumors and re-applying K-means clustering (K = 3). Tumors are from the 2013 TCGA sample set for uterine cancer and the heatmap depicts DDR gene expression from our curated list (131 genes). DDR gene expression groupings (Groups 1-6), identified by K-means clustering of DDR gene expression in Fig. 1A, are depicted on the heatmap.

**Supplemental Fig. 3. Expression of DDR genes in endometrial cancer with supervised clustering based on TCGA genomic subtypes.** Endometrial tumors (serous and endometrioid) clustered by TCGA genomic subtypes. DDR gene expression groupings (Groups 1-6) defined by K-means clustering based on DDR gene expression patterns identified in Fig. 1A are applied to the heatmap.

**Supplemental Fig. 4. Individual DDR genes predictive of overall survival in endometrial endometrioid carcinomas.** Kaplan-Meier curves for overall survival in endometrioid tumors with *POLE*-mutant tumors removed. Log-rank tests were used to infer statistical significance between high and low groups.

**Supplemental Fig. 5. Individual DDR genes predictive of progression free survival in endometrial endometrioid carcinomas.** Kaplan-Meier curves for overall survival in endometrioid tumors with *POLE*-mutant tumors removed. Log-rank tests were used to infer statistical significance between high and low groups.

**Supplemental Fig. 6. Identified DDR signatures and *POLQ* association are not generalizable to all HRD high tumors.** (A) Clustering of ESC, high grade ovarian serous carcinoma (HGOSC), and pancreatic ductal adenocarcinoma (PDAC) by curated DDR gene list (131 genes) (A) and at the DNA repair pathway level (B). ESC and HGOSC share similar histopathology and clinical features. (C) Wilcoxon’s rank sum test of HRD scores; graph is of PDAC, HGOSC, ESC, and high grade and low grade EEC in log2 scale. (D) Spearman’s rank correlation of *POLQ* expression and HRD score (in log2 scale) in HGSOC, ESC, and PDAC. (E) Spearman’s rank correlation of cosmic signature 3 vs. POLQ expression. ****p < 0.0001.

**Supplemental Fig. 7. Expression of DDR genes in endometrioid endometrial cancer with removing grade effect.** DDR gene expression in EEC. Clusters (Clusters 1-3) and groupings (Groups 1-6) defined by K-means clustering based on DDR gene expression patterns identified in Fig. 1A and 1F are applied to the heatmap.

**Supplemental Fig. 8. Expression of CTA genes based on TCGA genomic subtypes and HRD-high tumors.** Endometrial tumors (serous and endometrioid) clustered by TCGA genomic subtypes (A). DDR gene expression groupings (Groups 1-6) defined by K-means clustering based on DDR gene expression patterns identified in Fig. 1A are applied to the CTA gene expression heatmap. (B) CTA gene expression in HRD-high tumors.

**Supplemental Fig. 9. Validation and supporting data of CTAs in endometrial tumorigenesis.** Immunoblots probed for KIF2C (A). CEP55 (B), and NUF2 (C) showing knockdown efficiencies in HEC-1-B cells (A) and iHEuEC cells (B and C) 72 hours after transfection. (D) Immunoblots for NUF2 expression in iHEuEC EV and *KRAS^G12V^* cells with indicated siRNA, 72 after transfection. LE = long exposure and SE = short exposure. (E) Quantification of growth in HEC-1-B cells transfected with three different siRNA targeting KIF2C, +/- MK1775 (at indicated doses), analyzed by clonogenic survival assay. (F-G) Quantification of percentage of cells in different phases of cell cycle. HEC-1-B cells transfected with different siRNA targeting KIF2C +/- MK1775 (150 nM) treatment for 48h. Graph indicates percentage of G1, S, and G2/M population along with S-phase defects that can be visualized as collapsed S-phase populations. (F, G, and H) Two-way ANOVA, Tukey post-test, mean ± SD. *p < 0.05, **p < 0.005, ***p < 0.0005, ****p < 0.00005. ns = non-significant.

## References

1 Walens, A. et al. RNA-Based Classification of Homologous Recombination Deficiency in Racially Diverse Patients with Breast Cancer. Cancer Epidemiol Biomarkers Prev 31, 2136–2147, doi:10.1158/1055-9965.EPI-22-0590 (2022).

2 Parker, J. S. et al. Supervised risk predictor of breast cancer based on intrinsic subtypes. J Clin Oncol 27, 1160–1167, doi:10.1200/JCO.2008.18.1370 (2009).

3 Prat, A., Parker, J. S., Fan, C. & Perou, C. M. PAM50 assay and the three-gene model for identifying the major and clinically relevant molecular subtypes of breast cancer. Breast Cancer Res Treat 135, 301–306, doi:10.1007/s10549-012-2143-0 (2012).

4 Hamilton, C. A. et al. Uterine papillary serous and clear cell carcinomas predict for poorer survival compared to grade 3 endometrioid corpus cancers. British journal of cancer 94, 642–646, doi:10.1038/sj.bjc.6603012 (2006).

5 Clarke, M. A., Devesa, S. S., Harvey, S. V. & Wentzensen, N. Hysterectomy-Corrected Uterine Corpus Cancer Incidence Trends and Differences in Relative Survival Reveal Racial Disparities and Rising Rates of Nonendometrioid Cancers. J Clin Oncol 37, 1895–1908, doi:10.1200/JCO.19.00151 (2019).

6. Cancer Genome Atlas Research, N., et al. Integrated genomic characterization of endometrial carcinoma. Nature 497, 67-73, doi:10.1038/nature12113 (2013).

7 Dou, Y. et al. Proteogenomic Characterization of Endometrial Carcinoma. Cell 180, 729–748 e726, doi:10.1016/j.cell.2020.01.026 (2020).

8 Moxley, K. M. & McMeekin, D. S. Endometrial carcinoma: a review of chemotherapy, drug resistance, and the search for new agents. Oncologist 15, 1026–1033, doi:10.1634/theoncologist.2010-0087 (2010).

9 Whitehurst, A. W. Cause and consequence of cancer/testis antigen activation in cancer. Annu Rev Pharmacol Toxicol 54, 251–272, doi:10.1146/annurev-pharmtox-011112-140326 (2014).

10 Gao, Y. et al. A neomorphic cancer cell-specific role of MAGE-A4 in trans-lesion synthesis. Nature communications 7, 12105, doi:10.1038/ncomms12105 (2016).

11 Gao, Y., Tateishi, S. & Vaziri, C. Pathological Trans-Lesion Synthesis in Cancer. Cell Cycle, 0, doi:10.1080/15384101.2016.1214045 (2016).

12 Gao, Y. et al. The Cancer/Testes (CT) Antigen HORMAD1 promotes Homologous Recombinational DNA Repair and Radioresistance in Lung adenocarcinoma cells. Scientific reports 8, 15304, doi:10.1038/s41598-018-33601-w (2018).

13 Nichols, B. A. et al. HORMAD1 Is a Negative Prognostic Indicator in Lung Adenocarcinoma and Specifies Resistance to Oxidative and Genotoxic Stress. Cancer Res 78, 6196–6208, doi:10.1158/0008-5472.CAN-18-1377 (2018).

14 Jay, A., Reitz, D., Namekawa, S. H. & Heyer, W. D. Cancer testis antigens and genomic instability: More than immunology. DNA Repair (Amst) 108, 103214, doi:10.1016/j.dnarep.2021.103214 (2021).

15 Coll-de la Rubia, E., et al. Prognostic Biomarkers in Endometrial Cancer: A Systematic Review and Meta-Analysis. J Clin Med 9, doi:10.3390/jcm9061900 (2020).

16 Gates, E. J., Hirschfield, L., Matthews, R. P. & Yap, O. W. Body mass index as a prognostic factor in endometrioid adenocarcinoma of the endometrium. J Natl Med Assoc 98, 1814–1822 (2006).

17 Bowry, A., Kelly, R. D. W. & Petermann, E. Hypertranscription and replication stress in cancer. Trends Cancer 7, 863–877, doi:10.1016/j.trecan.2021.04.006 (2021).

18 Yang, Y. et al. DNA repair factor RAD18 and DNA polymerase Polkappa confer tolerance of oncogenic DNA replication stress. J Cell Biol, doi:10.1083/jcb.201702006 (2017).

19 Yang, Y. et al. Diverse roles of RAD18 and Y-family DNA polymerases in tumorigenesis. Cell Cycle, 1–11, doi:10.1080/15384101.2018.1456296 (2018).

20 Vaziri, C., Rogozin, I. B., Gu, Q., Wu, D. & Day, T. A. Unravelling roles of error-prone DNA polymerases in shaping cancer genomes. Oncogene 40, 6549–6565, doi:10.1038/s41388-021-02032-9 (2021).

21 Cong, K. & Cantor, S. B. Exploiting replication gaps for cancer therapy. Mol Cell 82, 2363–2369, doi:10.1016/j.molcel.2022.04.023 (2022).

22 Godinho, S. A. & Pellman, D. Causes and consequences of centrosome abnormalities in cancer. Philosophical transactions of the Royal Society of London 369, doi:10.1098/rstb.2013.0467 (2014).

23 Ogden, A., Rida, P. C. & Aneja, R. Prognostic value of CA20, a score based on centrosome amplification-associated genes, in breast tumors. Scientific reports 7, 262, doi:10.1038/s41598-017-00363-w (2017).

24 de Almeida, B. P., Vieira, A. F., Paredes, J., Bettencourt-Dias, M. & Barbosa-Morais, N. L. Pan-cancer association of a centrosome amplification gene expression signature with genomic alterations and clinical outcome. PLoS computational biology 15, e1006832, doi:10.1371/journal.pcbi.1006832 (2019).

25 McConechy, M. K. et al. Endometrial Carcinomas with POLE Exonuclease Domain Mutations Have a Favorable Prognosis. Clin Cancer Res 22, 2865–2873, doi:10.1158/1078-0432.CCR-15-2233 (2016).

26 Billingsley, C. C., Cohn, D. E., Mutch, D. G., Hade, E. M. & Goodfellow, P. J. Prognostic Significance of POLE Exonuclease Domain Mutations in High-Grade Endometrioid Endometrial Cancer on Survival and Recurrence: A Subanalysis. Int J Gynecol Cancer 26, 933–938, doi:10.1097/IGC.0000000000000681 (2016).

27 Leon-Castillo, A. et al. Clinicopathological and molecular characterisation of ’multiple-classifier’ endometrial carcinomas. The Journal of pathology 250, 312–322, doi:10.1002/path.5373 (2020).

28 Das Ghosh, D., et al. In-silico analysis of TCGA data showing multiple POLE-like favourable subgroups overlapping with TP53 mutated endometrial cancer: Implications for clinical practice in low and middle-income countries. Gynecol Oncol Rep 47, 101209, doi:10.1016/j.gore.2023.101209 (2023).

29 Rashid, S., Arafah, M. A. & Akhtar, M. The Many Faces of Serous Neoplasms and Related Lesions of the Female Pelvis: A Review. Adv Anat Pathol 29, 154–167, doi:10.1097/PAP.0000000000000334 (2022).

30 Ceccaldi, R. et al. Homologous-recombination-deficient tumours are dependent on Poltheta-mediated repair. Nature 518, 258–262, doi:10.1038/nature14184 (2015).

31 van Schendel, R., van Heteren, J., Welten, R. & Tijsterman, M. Genomic Scars Generated by Polymerase Theta Reveal the Versatile Mechanism of Alternative End-Joining. PLoS Genet 12, e1006368, doi:10.1371/journal.pgen.1006368 (2016).

32 Marquard, A. M. et al. Pan-cancer analysis of genomic scar signatures associated with homologous recombination deficiency suggests novel indications for existing cancer drugs. Biomark Res 3, 9, doi:10.1186/s40364-015-0033-4 (2015).

33 Kuramoto, H. et al. Establishment and characterization of human endometrial cancer cell lines. Annals of the New York Academy of Sciences 622, 402–421, doi:10.1111/j.1749-6632.1991.tb37884.x (1991).

34 Bakhoum, S. F., Thompson, S. L., Manning, A. L. & Compton, D. A. Genome stability is ensured by temporal control of kinetochore-microtubule dynamics. Nat Cell Biol 11, 27–35, doi:10.1038/ncb1809 (2009).

35 Liu, J. F. et al. Phase II Study of the WEE1 Inhibitor Adavosertib in Recurrent Uterine Serous Carcinoma. J Clin Oncol 39, 1531–1539, doi:10.1200/JCO.20.03167 (2021).

36 Madariaga, A. & Oza, A. M. Wee1 Inhibition in Recurrent Serous Uterine Cancer: Science Paving the Way in a Challenging Disease. J Clin Oncol 39, 1513–1517, doi:10.1200/JCO.21.00288 (2021).

37 Greene, B. L. et al. Ribonucleotide Reductases: Structure, Chemistry, and Metabolism Suggest New Therapeutic Targets. Annu Rev Biochem 89, 45–75, doi:10.1146/annurev-biochem-013118-111843 (2020).

38 Leman, A. R. & Noguchi, E. Local and global functions of Timeless and Tipin in replication fork protection. Cell Cycle 11, 3945–3955, doi:10.4161/cc.21989 (2012).

39 Reinhardt, H. C. & Yaffe, M. B. Kinases that control the cell cycle in response to DNA damage: Chk1, Chk2, and MK2. Curr Opin Cell Biol 21, 245–255 (2009).

40 Anand, J. et al. Roles of trans-lesion synthesis (TLS) DNA polymerases in tumorigenesis and cancer therapy. NAR Cancer 5, zcad005, doi:10.1093/narcan/zcad005 (2023).

41 Krais, J. J. et al. RNF168-mediated localization of BARD1 recruits the BRCA1-PALB2 complex to DNA damage. Nature communications 12, 5016, doi:10.1038/s41467-021-25346-4 (2021).

42 Garaycoechea, J. I. & Patel, K. J. Why does the bone marrow fail in Fanconi anemia? Blood 123, 26–34, doi:10.1182/blood-2013-09-427740 (2014).

43 Kotsantis, P., Petermann, E. & Boulton, S. J. Mechanisms of Oncogene-Induced Replication Stress: Jigsaw Falling into Place. Cancer discovery 8, 537–555, doi:10.1158/2159-8290.CD-17-1461 (2018).

44 Petropoulos, M., Champeris Tsaniras, S., Taraviras, S. & Lygerou, Z. Replication Licensing Aberrations, Replication Stress, and Genomic Instability. Trends Biochem Sci 44, 752–764, doi:10.1016/j.tibs.2019.03.011 (2019).

45 Bartkova, J. et al. Oncogene-induced senescence is part of the tumorigenesis barrier imposed by DNA damage checkpoints. Nature 444, 633–637, doi:10.1038/nature05268 (2006).

46 Godinho, S. A., Kwon, M. & Pellman, D. Centrosomes and cancer: how cancer cells divide with too many centrosomes. Cancer Metastasis Rev 28, 85–98, doi:10.1007/s10555-008-9163-6 (2009).

47 Levine, M. S. et al. Centrosome Amplification Is Sufficient to Promote Spontaneous Tumorigenesis in Mammals. Developmental cell 40, 313–322 e315, doi:10.1016/j.devcel.2016.12.022 (2017).

48 Fukasawa, K. Centrosome amplification, chromosome instability and cancer development. Cancer letters 230, 6–19, doi:10.1016/j.canlet.2004.12.028 (2005).

49 Godinho, S. A. et al. Oncogene-like induction of cellular invasion from centrosome amplification. Nature 510, 167–171, doi:10.1038/nature13277 (2014).

50 Passerini, V. et al. The presence of extra chromosomes leads to genomic instability. Nature communications 7, 10754, doi:10.1038/ncomms10754 (2016).

51 Sharma, S. S., Ma, L., Bagui, T. K., Forinash, K. D. & Pledger, W. J. A p27Kip1 mutant that does not inhibit CDK activity promotes centrosome amplification and micronucleation. Oncogene 31, 3989–3998, doi:10.1038/onc.2011.550 (2012).

52 Loffler, H., Lukas, J., Bartek, J. & Kramer, A. Structure meets function--centrosomes, genome maintenance and the DNA damage response. Exp Cell Res 312, 2633–2640, doi:10.1016/j.yexcr.2006.06.008 (2006).

53 Ganem, N. J. & Pellman, D. Linking abnormal mitosis to the acquisition of DNA damage. J Cell Biol 199, 871–881, doi:10.1083/jcb.201210040 (2012).

54 Yang, Z., Loncarek, J., Khodjakov, A. & Rieder, C. L. Extra centrosomes and/or chromosomes prolong mitosis in human cells. Nat Cell Biol 10, 748–751, doi:10.1038/ncb1738 (2008).

55 Hayashi, M. T. & Karlseder, J. DNA damage associated with mitosis and cytokinesis failure. Oncogene 32, 4593–4601, doi:10.1038/onc.2012.615 (2013).

56 Pavanello, L., Hall, B., Airhihen, B. & Winkler, G. S. The central region of CNOT1 and CNOT9 stimulates deadenylation by the Ccr4-Not nuclease module. Biochem J 475, 3437–3450, doi:10.1042/BCJ20180456 (2018).

57 Boussouar, F., Jamshidikia, M., Morozumi, Y., Rousseaux, S. & Khochbin, S. Malignant genome reprogramming by ATAD2. Biochim Biophys Acta 1829, 1010–1014, doi:10.1016/j.bbagrm.2013.06.003 (2013).

58 Salmon, E. D., Cimini, D., Cameron, L. A. & DeLuca, J. G. Merotelic kinetochores in mammalian tissue cells. Philosophical transactions of the Royal Society of London 360, 553–568, doi:10.1098/rstb.2004.1610 (2005).

59 Fujita, Y. et al. Priming of centromere for CENP-A recruitment by human hMis18alpha, hMis18beta, and M18BP1. Developmental cell 12, 17–30, doi:10.1016/j.devcel.2006.11.002 (2007).

60 Gaudet, S., Branton, D. & Lue, R. A. Characterization of PDZ-binding kinase, a mitotic kinase. Proc Natl Acad Sci U S A 97, 5167–5172, doi:10.1073/pnas.090102397 (2000).

61 Lan, W. & Cleveland, D. W. A chemical tool box defines mitotic and interphase roles for Mps1 kinase. J Cell Biol 190, 21–24, doi:10.1083/jcb.201006080 (2010).

62 Sinha, D., Duijf, P. H. G. & Khanna, K. K. Mitotic slippage: an old tale with a new twist. Cell Cycle 18, 7–15, doi:10.1080/15384101.2018.1559557 (2019).

63 Funabiki, H. & Wynne, D. J. Making an effective switch at the kinetochore by phosphorylation and dephosphorylation. Chromosoma 122, 135–158, doi:10.1007/s00412-013-0401-5 (2013).

64 Walczak, C. E. The Kin I kinesins are microtubule end-stimulated ATPases. Mol Cell 11, 286–288, doi:10.1016/s1097-2765(03)00067-4 (2003).

65 Cappell, K. M. et al. Multiple cancer testis antigens function to support tumor cell mitotic fidelity. Mol Cell Biol 32, 4131–4140, doi:10.1128/MCB.00686-12 (2012).

66 Weaver, B. A. How Taxol/paclitaxel kills cancer cells. Mol Biol Cell 25, 2677–2681, doi:10.1091/mbc.E14-04-0916 (2014).

67 Goldman, M. J. et al. Visualizing and interpreting cancer genomics data via the Xena platform. Nat Biotechnol 38, 675–678, doi:10.1038/s41587-020-0546-8 (2020).

68 Knijnenburg, T. A. et al. Genomic and Molecular Landscape of DNA Damage Repair Deficiency across The Cancer Genome Atlas. Cell reports 23, 239–254 e236, doi:10.1016/j.celrep.2018.03.076 (2018).

69 Konstantinopoulos, P. A. et al. A Replication stress biomarker is associated with response to gemcitabine versus combined gemcitabine and ATR inhibitor therapy in ovarian cancer. Nature communications 12, 5574, doi:10.1038/s41467-021-25904-w (2021).

70 Gu, Z., Eils, R. & Schlesner, M. Complex heatmaps reveal patterns and correlations in multidimensional genomic data. *Bioinformatics (Oxford*, England) 32, 2847–2849, doi:10.1093/bioinformatics/btw313 (2016).

