## Supplemental Figure Legends for "The DNA Damage Response (DDR) landscape of endometrial cancer defines discrete disease subtypes and reveals therapeutic opportunities"

### Supplemental Fig. 1

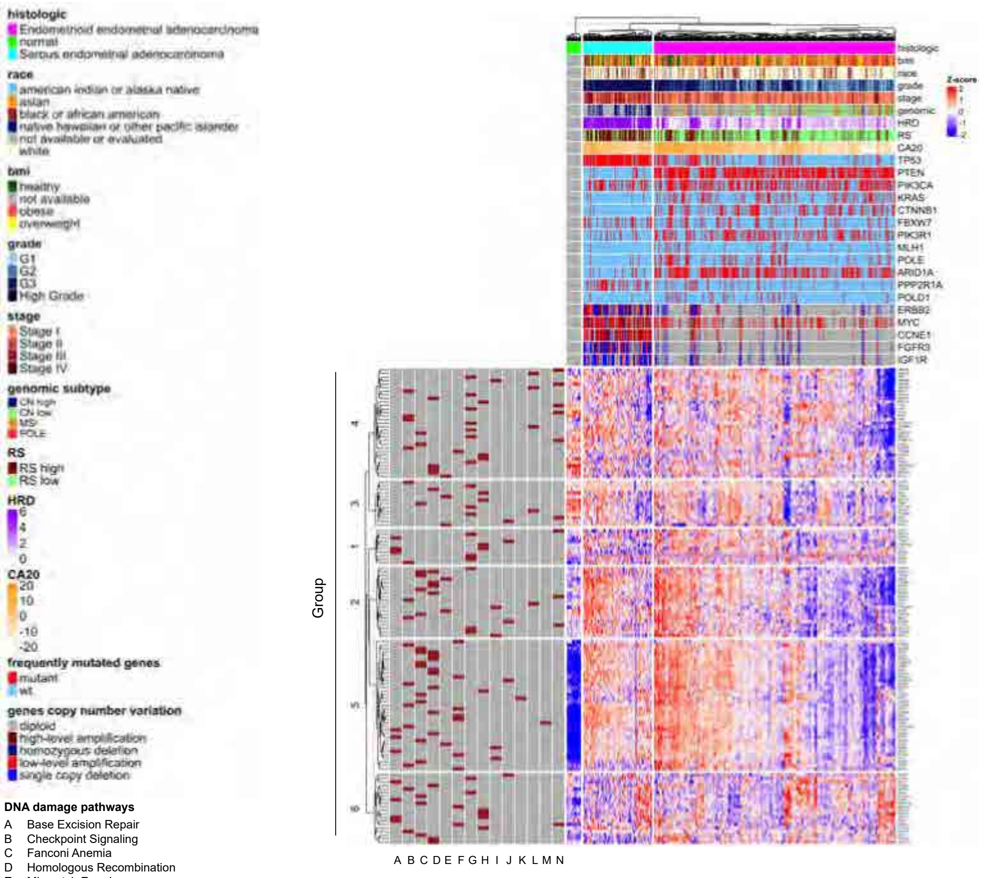


**DNA damage pathways**

A Base Excision Repair
B Checkpoint Signaling
C Fanconi Anemia
D Homologous Recombination
E Mismatch Repair
F Mitosis/Spindle Assembly Checkpoint
G NHEJ Pathway
H Nucleotide Excision Repair
I Nucleotide Metabolism
J Poly ADP Ribose Polymerases
K Repair/Replication DNA Polymerases
L Template Switch
M Theta-Mediated End Joining
N Trans-Lesion Synthesis

#### Supplemental Fig. 2

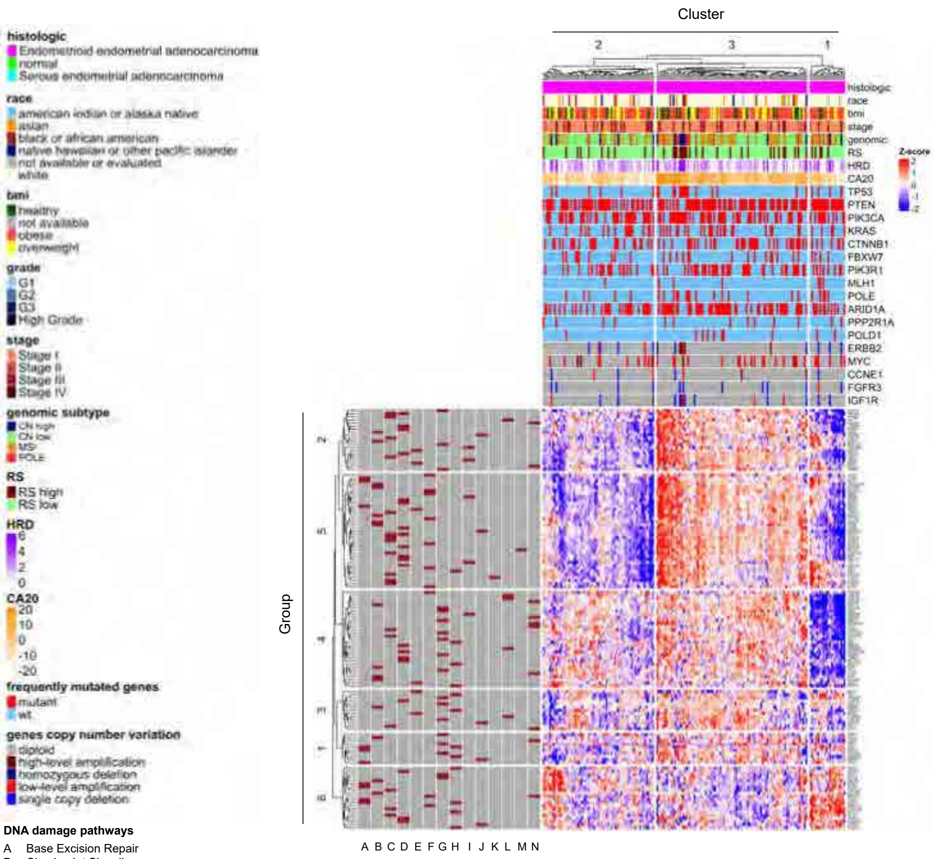


**DNA damage pathways**

A Base Excision Repair
B Checkpoint Signaling
C Fanconi Anemia
D Homologous Recombination
E Mismatch Repair
F Mitosis/Spindle Assembly Checkpoint
G NHEJ Pathway
H Nucleotide Excision Repair
I Nucleotide Metabolism
J Poly ADP Ribose Polymerases
K Repair/Replication DNA Polymerases
L Template Switch
M Theta-Mediated End Joining
N Trans-Lesion Synthesis

**Supplemental Fig. 3**

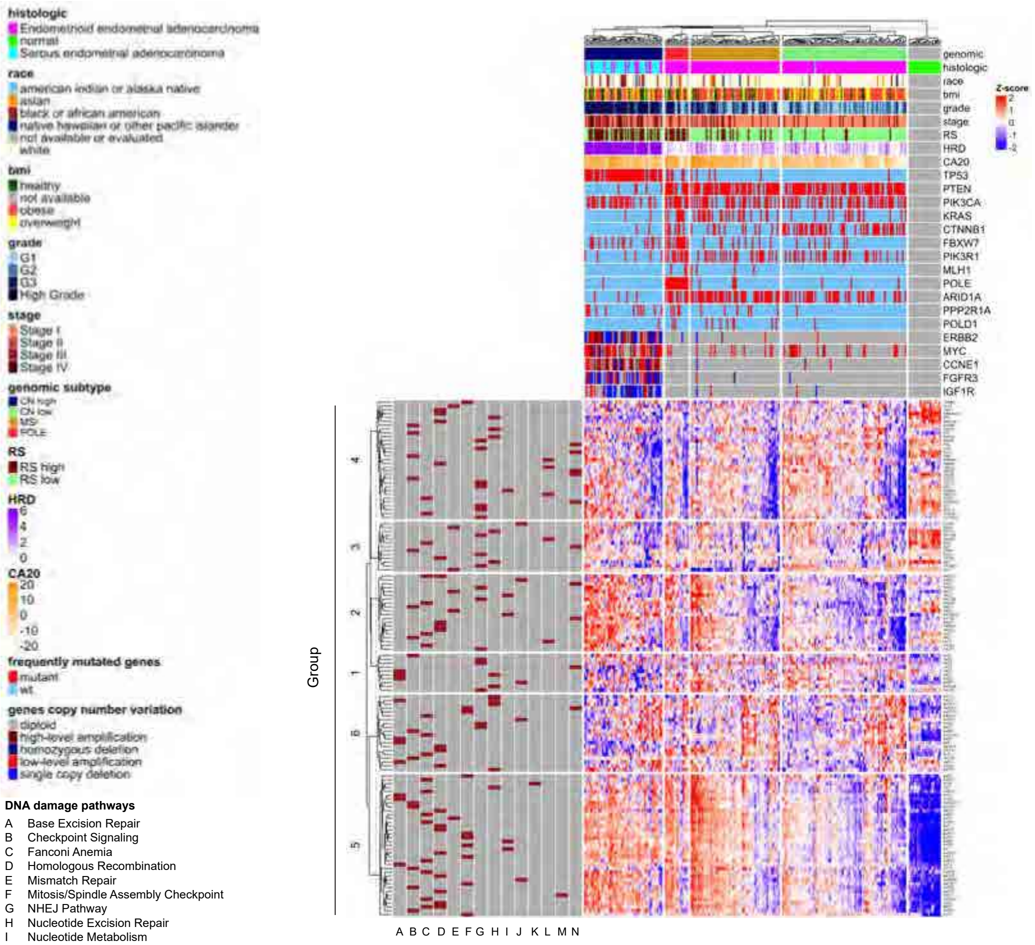


**DNA damage pathways**

A Base Excision Repair
B Checkpoint Signaling
C Fanconi Anemia
D Homologous Recombination
E Mismatch Repair
F Mitosis/Spindle Assembly Checkpoint
G NHEJ Pathway
H Nucleotide Excision Repair
I Nucleotide Metabolism
J Poly ADP Ribose Polymerases
K Repair/Replication DNA Polymerases
L Template Switch
M Theta-Mediated End Joining
N Trans-Lesion Synthesis

**Supplemental Fig. 4**

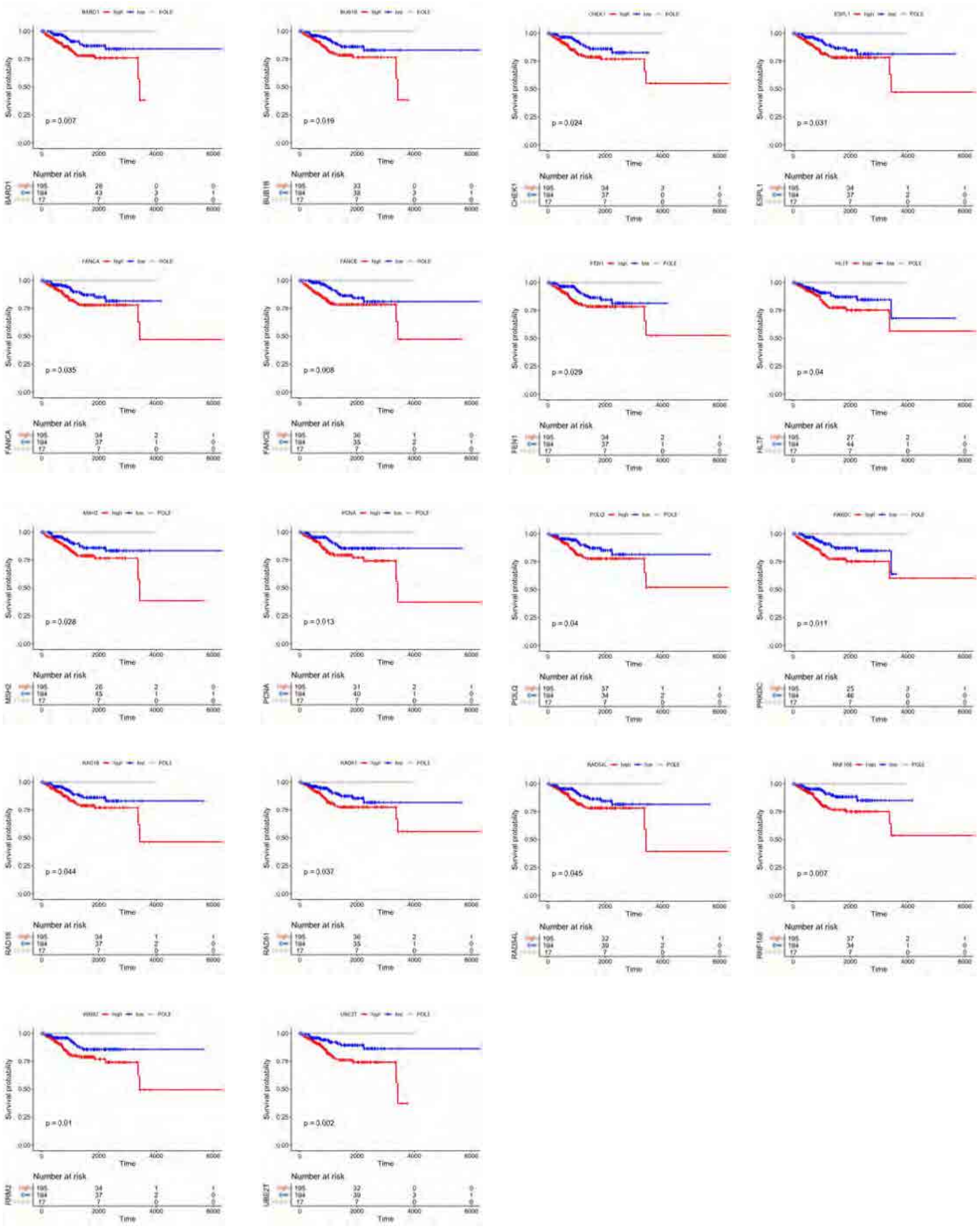

**Supplemental Fig. 5**

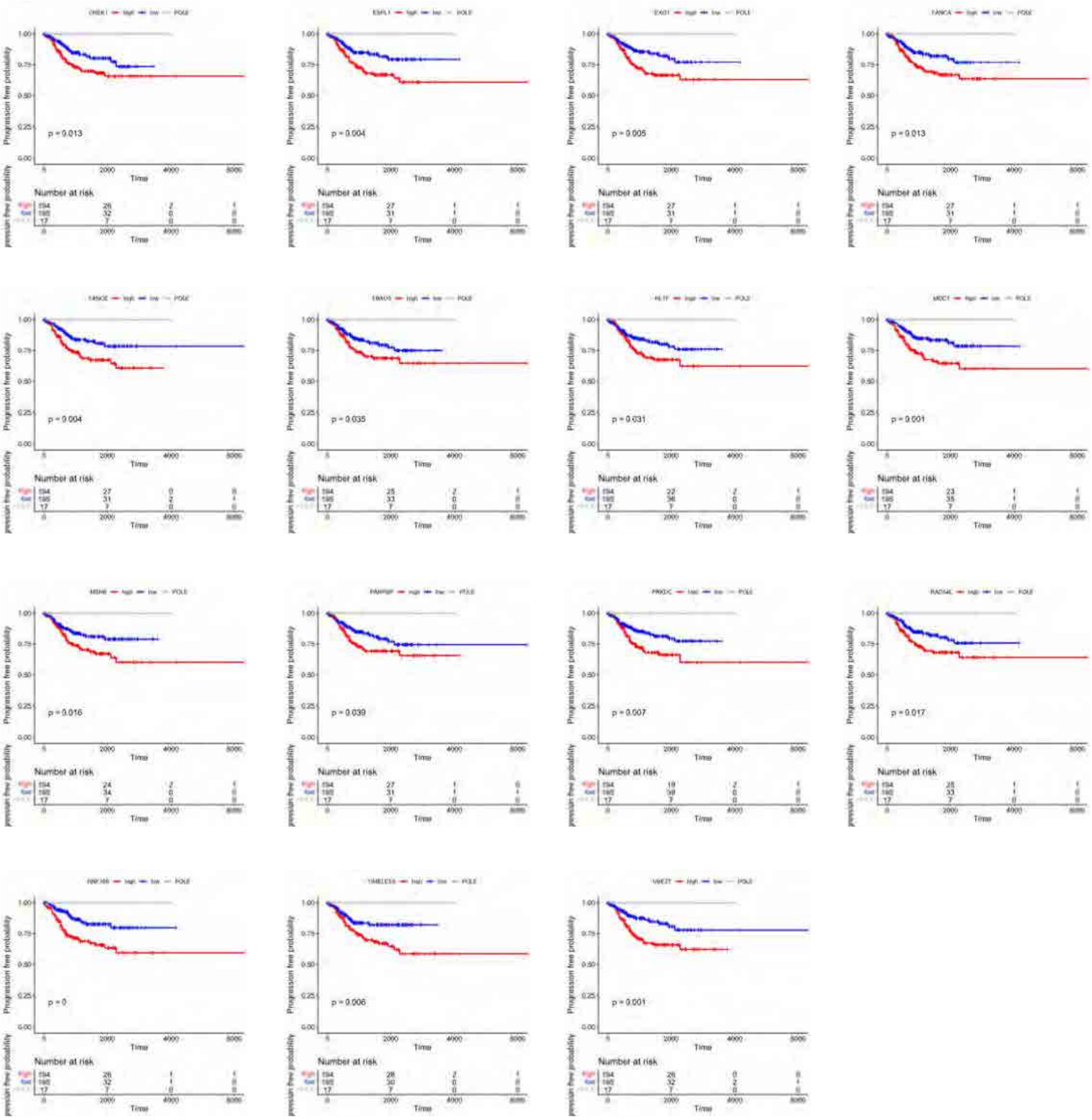

#### Supplemental Fig. 6

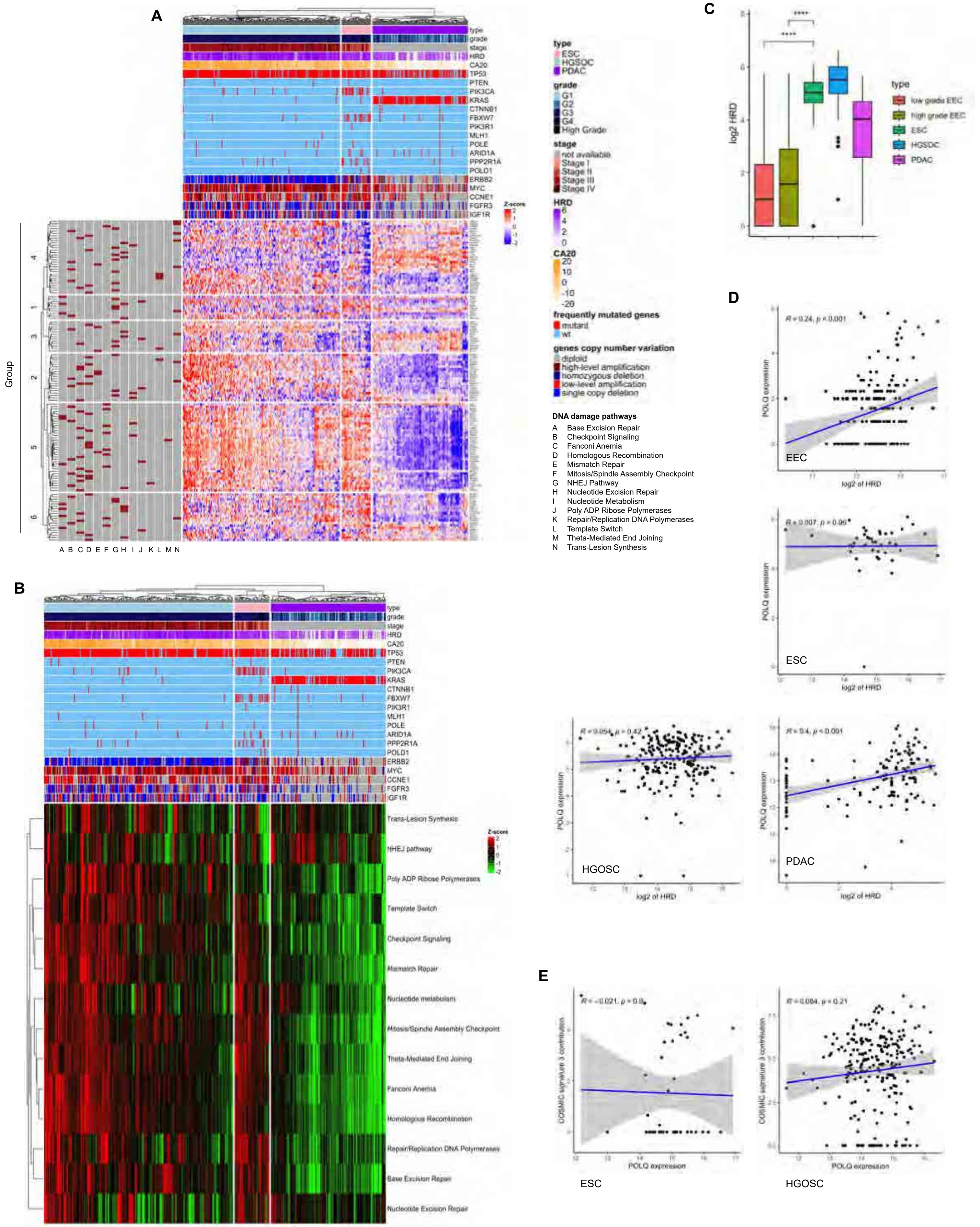

**Supplemental Fig. 7**

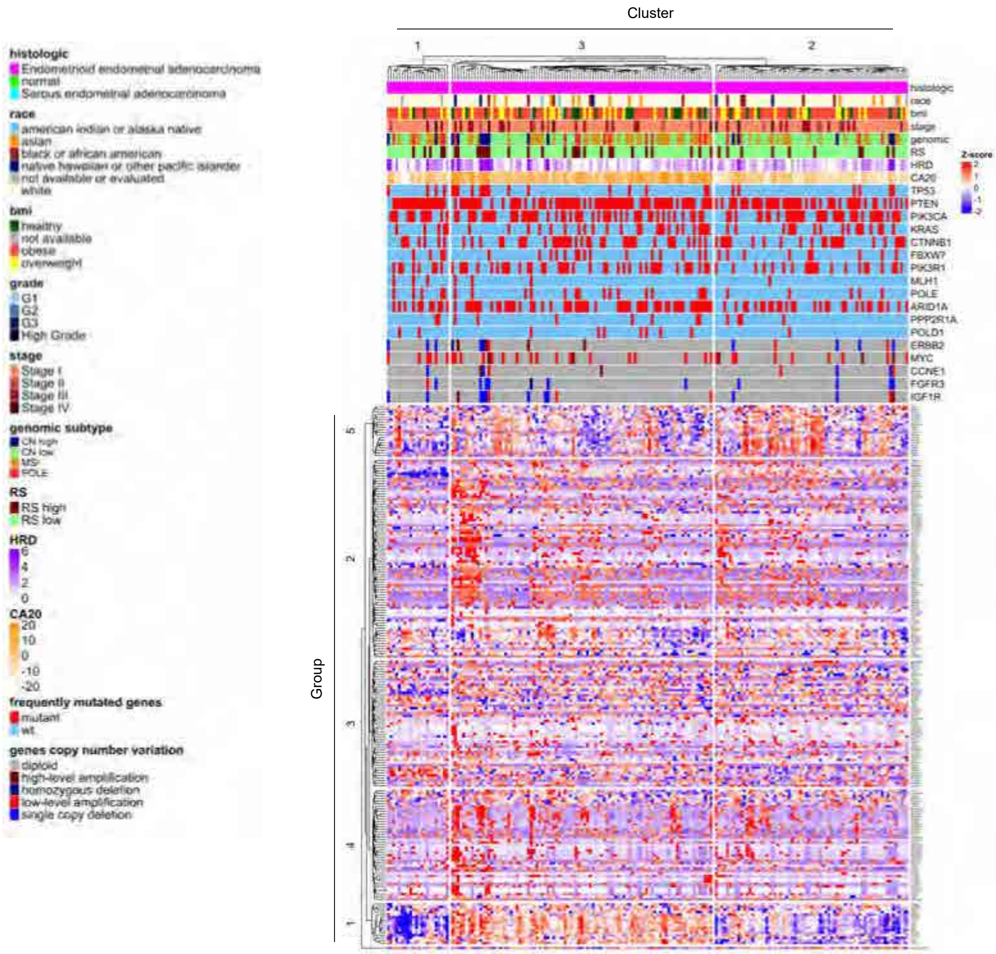

**Supplemental Fig. 8**

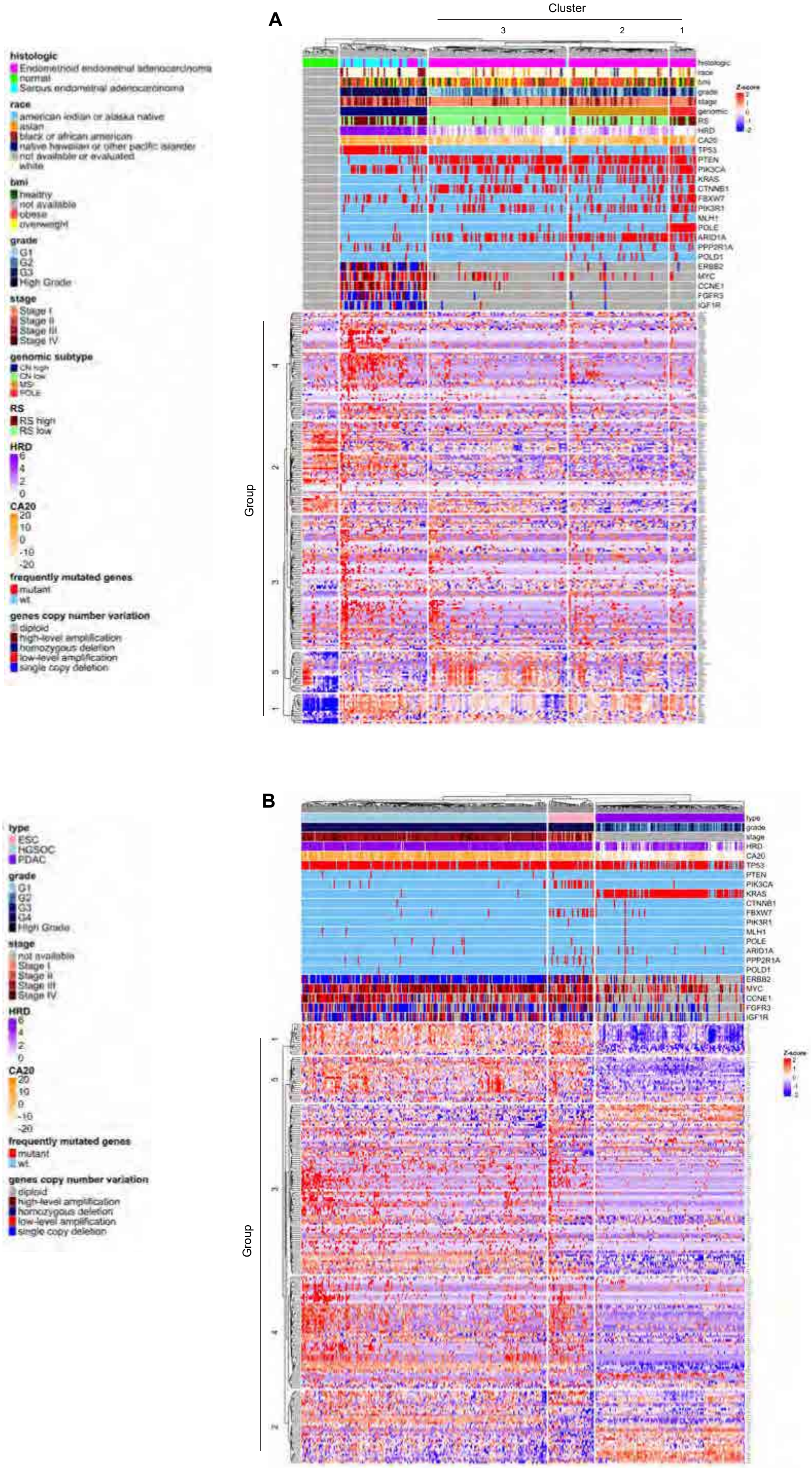

**Supplemental Fig. 9**

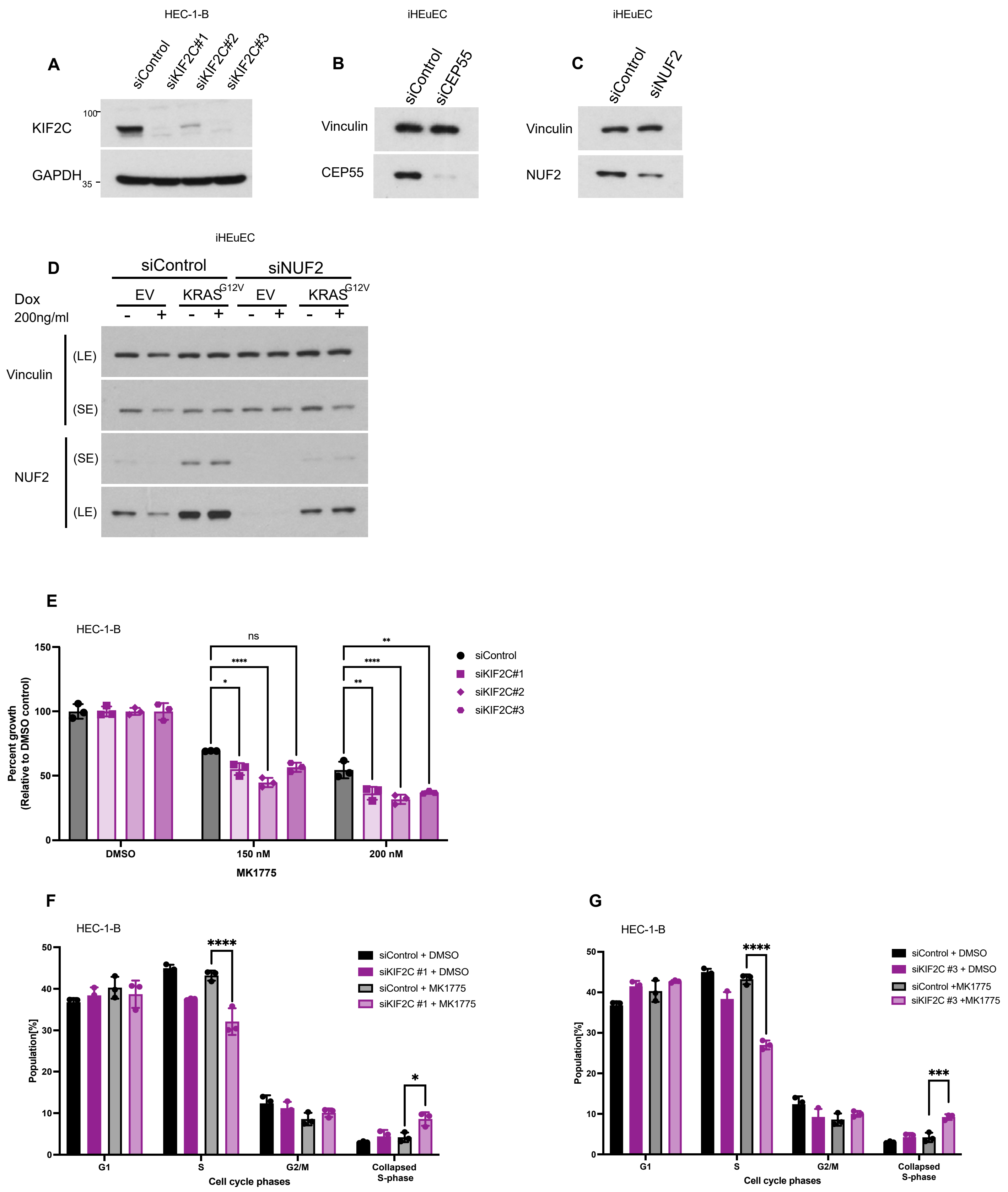
